# Muscle-secreted G-CSF as a metabolic niche factor ameliorates loss of muscle stem cell in aged mice

**DOI:** 10.1101/488874

**Authors:** Hu Li, Qian Chen, Changyin Li, Ran Zhong, Yixia Zhao, Qianying Zhang, Weimin Tong, Dahai Zhu, Yong Zhang

## Abstract

Function and number of muscle stem cells (satellite cells, SCs) declines with muscle aging. Although SCs are heterogeneous and different subpopulations have been identified, it remains unknown if a specific subpopulation of muscle SCs selectively decreases during aging. Here, we find Pax7^Hi^ cells are dramatically reduced in aged mice and this aged-dependent loss of Pax7^Hi^ cells is metabolically mediated by myofiber-secreted granulocyte-colony stimulating factor G-CSF as the Pax7^Hi^ SCs are replenished by exercise-induced G-CSF in aged mice. Mechanistically, we show that transcription of G-CSF (*Csf3*) gene in myofibers is regulated by MyoD in a metabolism-dependent manner and the myofibers-secreted G-CSF acts as a metabolic niche factor required for establishing and maintaining the Pax7^Hi^ SC subpopulation in adult and physiological aged mice by promoting the asymmetric division of Pax7^Hi^ and Pax7^Mi^ SCs. Together, our findings uncover a metabolic niche role of muscle metabolism in regulating Pax7 SC heterogeneity in mice.

**Highlights:**

1. Single cell RNA-seq unveils Pax7^Hi^ and Pax7^Lo^ cells are two distinct subpopulations.
2. Pax7^Hi^ SCs are enriched in glycolytic fibers and reduced in aging muscle.
3. Metabolic niche factor G-CSF is required for regulating dynamic change of Pax7 SCs.
4. G-CSF replenishes Pax7^Hi^ cells by stimulating asymmetric division of Pax7^Mi^ cells.

## Introduction

Reduced tissue regenerative potential is one of the general hallmarks in mammalian aging (Rando, 2006) and decline in the number and function of adult stem cells is the major causes that contribute to the failure of regeneration in several adult tissues during aging (Conboy, Conboy et al., 2003, Molofsky, Slutsky et al., 2006, Nishimura, Granter et al., 2005, Rossi, Mash et al., 2005). However, the molecular mechanisms underlying this age-dependent loss of adult stem cells in tissue regeneration are largely unknown. In adult skeletal muscle, muscle stem cells, also known as satellite cells (SCs), reside in a quiescent state between the basal lamina and the muscle fiber sarcolemma. SCs are responsible for postnatal muscle growth and regeneration after injury. SCs are also heterogeneous where different subpopulations have unique features of self-renewal, proliferation, and differentiation during regeneration (Kuang, Kuroda et al., 2007, Rocheteau, Gayraud-Morel et al., 2012, Wilson, Laurenti et al., 2008). Pax7, a transcriptional factor, plays critical roles in regulating SC functions during development and regeneration (Sambasivan, Yao et al., 2011, Seale, Sabourin et al., 2000, von Maltzahn, Jones et al., 2013). It has been recently reported that Pax7-positive SCs (Pax7 SCs) are heterogeneous including Pax7^Hi^ and Pax7^Lo^ subpopulations (Rocheteau et al., 2012, Wu et al. 2015). However, little is known regarding the functional establishment and maintenance of the heterogeneity of Pax7 SCs during development. During aging, the decline in number and function of Pax7 SCs is attributable to the loss of skeletal muscle mass and strength as well as the decreased regenerative capacity. More intriguingly, it remains unclear if a specific subpopulation of Pax7 SCs decreases during aging and what are the dynamics of this heterogeneity in aged mice.

The microenvironment, or niche contributes significantly to the behaviors of adult stem cells, as first reported for germ stem cell niche of the *Drosophila* ovary (Xie & Spradling, 2000) and the hematopoietic stem cell niche in mammal (Schofield, 1978). However, little is known about which niche components are required to regulate the heterogeneity of adult stem cells. The identification of niche factors will help to elucidate the molecular mechanisms underlying the establishment and maintenance of adult stem cell heterogeneity during development and physiological aging. In skeletal muscle, Pax7 SCs were directly attached with two major types of muscle fibers which are defined based on their metabolic capacity: slow-twitch oxidative fibers and fast-twitch glycolytic fibers (Schiaffino & Reggiani, 2011). Interestingly, there is a link between the SC numbers/function and fiber metabolism, more SC cells on slow-twitch oxidative fibers than that on fast-twitch glycolytic fibers (Collins, Olsen et al., 2005, Feldman & Stockdale, 1991, Lagord, Soulet et al., 1998). Further, decline of SC numbers and function is correlated with fiber type switch from glycolytic fast-twitch to oxidative slow-twitch fiber during aging. These observations imply a possible effect of fiber metabolism on SC function and behavior during aging.

Skeletal muscle is a major secretory organ, and muscle fibers express and secrete various factors (e.g., IL-6 and FGF-2) that regulate skeletal muscle growth and regeneration in autocrine, paracrine, or endocrine manners (Pedersen & Febbraio, 2008, Pedersen & Febbraio, 2012). Given that muscle fibers exhibit metabolic heterogeneity, secrete factors that have paracrine function, and exhibit intimate contact with Pax7 SCs, we hypothesized that muscle fibers function as a metabolic niche for skeletal muscle SCs by supplying requisite factors that in turn regulate the heterogeneity of Pax7 SCs during development and aging in mice. In the present study, we tested this hypothesis using several experimental approaches. First, using single cell RNA sequencing, we demonstrate that Pax7^Hi^ and Pax7^Lo^ cells are two distinct subpopulations of satellite cells. More significantly, we uncover that the number of Pax7^Hi^ subpopulation satellite cells is significantly reduced in aged mice. Mechanistically, we reveal that altered heterogeneity of Pax7 SCs is regulated by myofiber-secreted granulocyte-colony stimulating factor (G-CSF), which is metabolically regulated by MyoD in myofibers and in turn interacts with its receptor, G-CSFR, on Pax7 SCs. This interaction is required for establishing and maintaining the Pax7^Hi^ SC subpopulation in adult and physiological aged mice by promoting the asymmetric division of Pax7^Hi^ and Pax7^Mi^ SCs.

## Results

### Characterization of Pax7^Hi^ and Pax7^Lo^ SCs by single-cell RNA sequencing

Quiescent Pax7^Hi^ and Pax7^Lo^ cells isolated by FACS based on levels of GFP (Pax7) from tibialis anterior (TA) muscle of *Pax7-nGFP* mice (Fig EV1A-D) were subjected to single cell RNA sequencing (scRNA-Seq). We profiled 1,243 Pax7^Hi^ cells and 3,960 Pax7^Lo^ cells. The typical number of detectable genes ranged approximately from 1000 to 2000 genes in individual cells. Unsupervised hierarchal clustering analysis with the single-cell RNA transcriptome indicated that quiescent Pax7^Hi^ and Pax7^Lo^ cells belonged to two distinctly clustered subpopulations (Fig 1A) as indicated with quiescent marker Vcam1 (Fig EV1E). Transcriptome comparisons between Pax7^Hi^ and Pax7^Lo^ subpopulations identified 428 differentially expression genes (LogFC>0.25), which exhibit distinct gene signatures (Fig 1B). Furthermore, GO-enriched analysis of the differentially expressed genes between those two subpopulations consistently validated the previously described features (Fig EV1F-H). Genes related to stemness were highly expressed in the Pax7^Hi^ subpopulation and genes related to myogenic differentiation were highly expressed in the Pax7^Lo^ subpopulation (Fig 1C). Additionally, we found that Pax7^Hi^ cells expressed high levels of mitochondrial genes (Figs 1D and EV1I), suggesting that Pax7^Hi^ cells were adapted to oxidative metabolism. Finally, several molecular markers for either Pax7^Hi^ or Pax7^Lo^ cells were identified in this study. *Ptprb*, *Pvalb*, *Acta1*, *Hbb-bt* are for Pax7^Hi^ cells and *Cdh15*, *Rcan2*, *Rps28*, *Acta2* for Pax7^Lo^ cells (Fig 1E and F). The expression patterns of these genes were validated by real-time PCR (Fig EV1J). Together, the high resolution analysis using single-cell RNA sequencing provides evidence that Pax7^Hi^ and Pax7^Lo^ cells represent two distinct subpopulations in mice. Therefore, the Pax7^Hi^ and Pax7^Lo^ cells used in the following experiments were FACS-sorted based on the levels of Pax7 expression as previously reported (Rocheteau et al., 2012, Wu et al., 2015).

**Fig 1.**
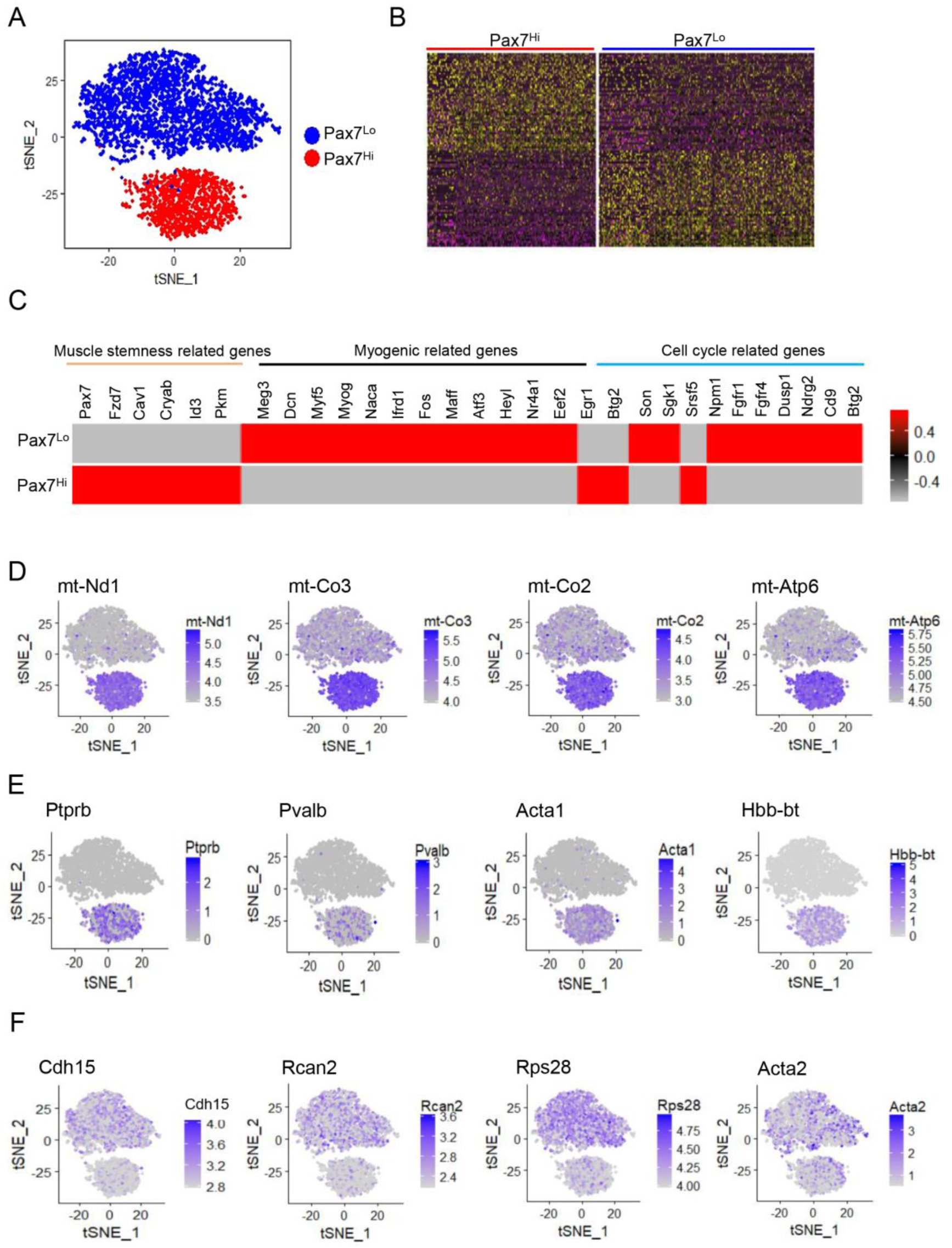
Transcriptional profile of Pax7^Hi^ and Pax7^Lo^ cells by single cell RNA seq. A Two-dimensional (2D) visualization of single-cell profiles inferred from RNA-seq data for Pax7^Hi^ and Pax7^Lo^ cells sorted from the TA muscle of young (3-month-old) *Pax7-nGFP* mice. Each point is a single cell colored by cluster assignment. B Heatmaps of normalized genes show Pax7^Hi^ and Pax7^Lo^ by top genes (columns) for individual cells (rows). C Differentially expressed genes between Pax7^Hi^ and Pax7^Lo^ cells in heatmap view. Genes were labeled with the molecular function, as indicated. D Expression patterns of *mt-Nd1*, *mt-Co3*, *mt-Co2* and *mt-Atp6* were visualized by t-SNE plots. E Top unique expressed genes in Pax7^Hi^ cells were visualized by t-SNE plots. F Top unique expressed genes in Pax7^Lo^ cells were visualized by t-SNE plots.

### Pax7^Hi^ cells are significantly reduced in aged mice

Given that the number and functionality of Pax7 SCs decline with age and Pax7^Hi^ cells with more stem-like properties represent a reversible dormant stem cell state and generate distinct daughter cell fates by asymmetrically segregating template DNA during muscle regeneration. We assessed whether the percentage of Pax7^Hi^ cells was altered in aged mice. Satellite cells were sorted from TA muscle of young and aged Pax7-nGFP mice, respectively. FACS profiling revealed that the percentage of Pax7^Hi^ SCs was severely reduced in the TA muscle fibers of aged mice compared to young mice (Fig2 A-C). Consistent with this, we observed low levels of *Pax7* and stemness-related genes *CD34*, but high levels of myogenic differentiation-related genes (*MyoD* and *MyoG*) in Pax7 SCs freshly isolated from aged TA muscle fibers versus young TA muscle fibers (Fig 2D). We also examined expression of newly identified genes as markers for either Pax7^Hi^ or Pax7^Lo^ cells, respectively. We found lower levels of Pax7^Hi^ marker genes expression (*mt-Nd1*, *mt-Co2* and *mt-Co3*) but higher levels of Pax7^Lo^ marker genes expression (*Rcan2* and *Cdh15*) in Pax7 SCs freshly isolated from aged TA muscle fibers versus young TA muscle fibers (Fig 2E). Consistently, we observed that the activation of Pax7 SCs was significantly accelerated in the cells freshly isolated from aged TA muscle fibers versus the young TA muscle fibers (Fig 2F and G).Taken together, these results demonstrate that Pax7^Hi^ cells were significant reduced in aged mice.

**Fig 2.**
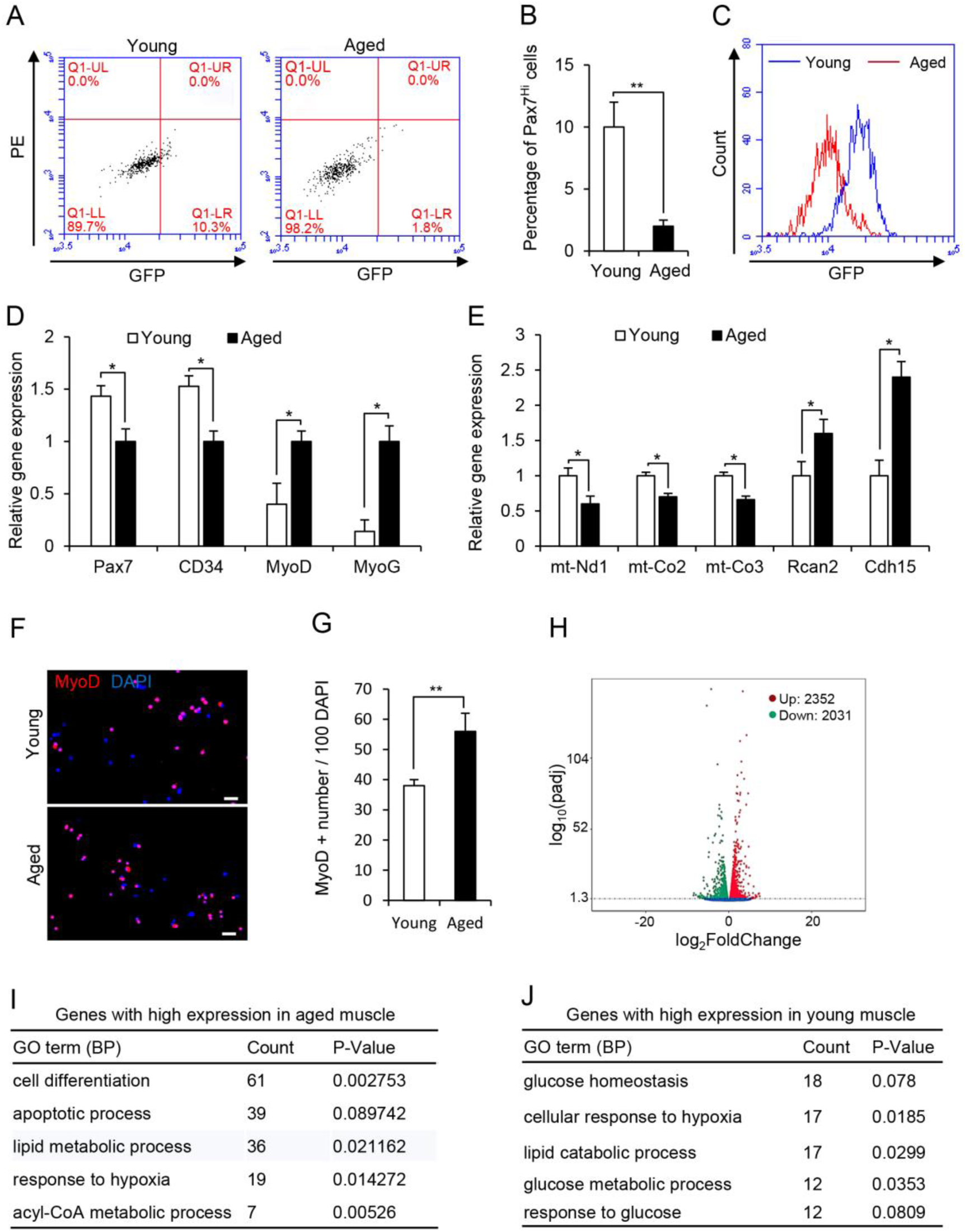
Reduced Pax7^Hi^ cells in aged mice. A Representative FACS profiles of Pax7 SCs from the TA muscles of aged (18-month-old) and young (3-month-old) *Pax7-nGFP* mice. B The percentages of Pax7^Hi^ SCs in (A) were calculated. *n* = 5 for each group. *p< 0.05. Unpaired two-sided t-test. C Representative overall profiles of the Pax7 SCs described in (A). D Relative expressions of molecular markers for stemness and differentiation in FACS-resolved Pax7 SCs from the TA muscles of aged and young *Pax7-nGFP* mice, as determined by qRT-PCR. *n* = 5. *p< 0.05. Unpaired two-sided t-test. E Relative expression of molecular markers for Pax7^Hi^ cells and Pax7^Lo^ cells in FACS-resolved Pax7 SCs from the TA muscles of aged and young *Pax7-nGFP* mice, as determined by qRT-PCR. *n* = 5. *p< 0.05. Unpaired two-sided t-test. F Representative images of MyoD immunostaining (red) for FACS-resolved Pax7 SCs cultured in growth medium (GM) for 18 hr. DAPI was used to visualize nuclei (blue). Scale bar represents 20 µm. G The percentages of MyoD-positive cells in (F) were calculated from 3 independent experiments. *p< 0.05. Unpaired two-sided t-test. H Volcano plot displayed the differentially expressed genes between aged and young TA muscles. Each point represents the log2FoldChange and log10(padj) from three independent biological replicates. Red is upregulated genes in aged muscle compared to young one. Green is downregulated genes in aged muscle compared to young one. Blue represents genes with no change between aged and young muscle. I,J Gene ontology (GO) analyses of the differentially expressed genes between aged and young TA muscle were shown as biological process (BP). The enriched GO terms for genes with high expression in aged or young TA muscle were shown in (I) and (J), respectively. “Count” stand for the number of genes enriched in the indicated GO term. The differentially expressed genes between aged and young TA muscle were identified with cutoff (fold change > 1.5, padj< 0.05).

### Glycolytic metabolism of myofiber is required for the maintenance of Pax7^Hi^ SC

To identify niche components in skeletal muscle that might be required for the maintenance of Pax7^Hi^ cells in aged mice, we analyzed the transcriptomes of aged TA muscle and young TA muscle by RNA sequencing. RNA-seq results from aged TA muscle and young TA muscle identified a large number of differentially regulated genes (Fig 2H). Gene ontology (GO) analyses of biological processes highlighted the changes in a number of genes encoding metabolic regulators in the transition from young to aged muscle (Fig 2I and J). The expression of genes corresponding to proteins that regulate lipid metabolic processes was upregulated (Figs 2I and EV2A), whereas that of genes regulating glucose metabolic processes was downregulated in aged TA muscle (Figs 2J and EV 2B and C).To further confirm the metabolic shift during aging, we performed histochemical staining for α-glycerophosphate dehydrogenase (α-GPDH) and succinate dehydrogenase (SDH), which are enriched in glycolytic and oxidative myofibers, respectively. Aged TA muscle had higher SDH and lower α-GPDH enzymatic activities compared to young TA muscles (Fig EV2D). Altogether, these results indicate that muscle metabolism was shifted from a glycolytic to oxidative state during aging as previously reported (Holloszy, Chen et al., 1991).

Since skeletal muscle composed of glycolytic myofiber and oxidative myofiber and SCs were directly attached with muscle fibers, we reasoned that metabolism of muscle fibers as a metabolic niche regulating the heterogeneity of Pax7 SC cells. To this end, we examined the distributions of the Pax7^Hi^ subpopulations in TA (predominantly glycolytic) and soleus (Sol, mainly oxidative) muscle fibers from the same individual *Pax7-nGFP* reporter mice. Interestingly, the percentage of Pax7^Hi^ SCs sorted from glycolytic TA muscle fibers was significantly higher than that sorted from oxidative Sol muscle fibers of the same animal (Fig 3A-C), indicating that the Pax7^Hi^ SC subpopulation was enriched in glycolytic muscle fibers of these mice. Consistent with this, we observed higher levels of *Pax7* and stemness-related genes (*CXCR4* and *CD34*) but lower levels of myogenic differentiation-related genes (*MyoG*) in Pax7 SCs freshly isolated from TA muscle fibers versus Sol muscle fibers of the same mice (Fig 3D). We also examined expression of newly identified genes as markers for either Pax7^Hi^ or Pax7^Lo^ cells, respectively. We found higher levels of Pax7^Hi^ marker genes expression (*mt-Nd1*, *mt-Co2*, *mt-Co3*, *Ptprb* and *Hbb-bt*) but lower levels of Pax7^Lo^ marker genes expression (*Rcan2* and *Cdh15*) in Pax7 SCs freshly isolated from TA muscle fibers versus Sol muscle fibers of the same animal (Fig 3E). Consistently, we observed that the activation of Pax7 SCs was significantly delayed in the cells freshly isolated from TA muscle fibers versus the corresponding Sol muscle fibers (Fig 3F and G). To rule out the possibility that enrichment of Pax7^Hi^ SCs in glycolytic TA muscle fibers is mediated by fiber type per se, we examined correlation between fiber type/fiber metabolism and the heterogeneity of Pax7 SC cells in TA and Sol muscle at 3 and 10 weeks, respectively. We found that the fiber type had been remarkably different between TA and Sol muscle of 3 weeks mice (Fig EV3A and B), but the significant difference in muscle metabolism was occurred till 10 weeks (Fig EV3C-E). Interestingly, the significant difference in the percentage of Pax7^Hi^ cells between TA and Sol muscle was only observed at 10 weeks but not at 3 weeks (Fig 3H), indicating that metabolic features of myofibers rather than muscle contraction associated with establishment of Pax7^Hi^ subpopulations in TA muscle. Our results show for the first time that Pax7 SCs are remarkably heterogeneous between glycolytic and oxidative muscle fibers. This heterogeneity may be related to the metabolic activity of these muscle fibers under normal physiological conditions.

**Fig 3.**
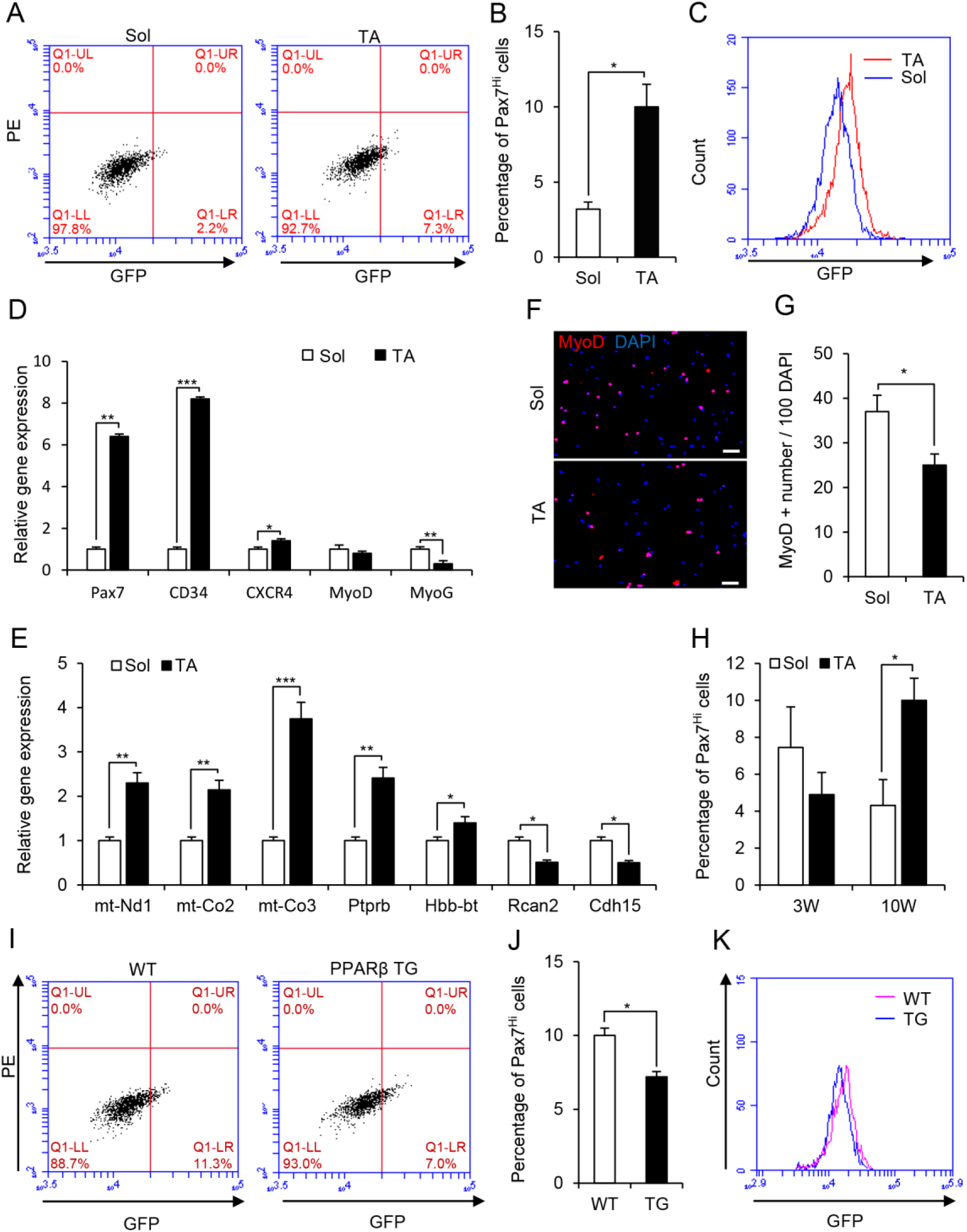
Glycolytic metabolism of myofiber associates with Pax7^Hi^ SC subpopulation. A Representative FACS profiles of Pax7 SCs sorted from Sol or TA muscles of *Pax7-nGFP* mice. GFP, green fluorescence protein (488 channel); PE, phycoerythrin (594 channel). B Average percentages of Pax7^Hi^ SC subpopulations obtained from 3 independent FACS experiments, performed as described in panel (A). *p< 0.05. Unpaired two-sided t-test. C Representative FACS profiles of total Pax7 SCs sorted from the Sol or TA muscles of *Pax7-nGFP* mice. GFP, green fluorescence protein (488 channel). D Relative expression of molecular markers for stemness and differentiation in FACS-resolved Pax7 SCs from the TA and Sol muscles of *Pax7-nGFP* mice, as determined by qRT-PCR. *n* = 5. *p< 0.05. **p< 0.01. ***p< 0.001. Unpaired two-sided t-test. E Relative expression of molecular markers in Pax7^Hi^ cells and Pax7^Lo^ cells of FACS-resolved Pax7 SCs from the TA and Sol muscles of *Pax7-nGFP* mice, as determined by qRT-PCR. *n* = 5. *p< 0.05. **p< 0.01. ***p< 0.001. Unpaired two-sided t-test. F Representative images of MyoD immunostaining (red) for FACS-resolved Pax7 SCs cultured in growth medium (GM) for 18 hr. DAPI was used to visualize nuclei (blue). Scale bar represents 20 µm. G The percentages of MyoD-positive cells in (F) were calculated from three independent experiments. H Average percentages of Pax7^Hi^ SC subpopulations in TA and Sol muscle from 3 weeks and 10 weeks *Pax7-nGFP* mice. Data were obtained from 3 independent FACS experiments. *p< 0.05. Unpaired two-sided t-test. I Representative FACS profiles of total Pax7 SCs sorted from the TA muscles of *Pax7-nGFP;MCK-PPARβ* transgenic (TG) and *Pax7-nGFP* wild-type (WT) littermates. GFP, green fluorescence protein (488 channel). J The percentages of Pax7^Hi^ SCs in (I) were calculated from 3 independent experiments. *p< 0.05. Unpaired two-sided t-test. K Representative FACS profiles of total Pax7 SCs sorted from the TA muscles of *Pax7-nGFP;MCK-PPARβ* transgenic (TG) and *Pax7-nGFP* wild-type (WT) littermates. GFP, green fluorescence protein (488 channel). *p< 0.05. **p< 0.01. ***p< 0.001. Unpaired two-sided t-test.

To further confirm the causal effects of fiber metabolism on Pax7 SC heterogeneity in mice, we examined the distribution of the Pax7^Hi^ SC subpopulation in the TA muscle of the well-characterized *PPARβ* transgenic (TG) mice, which exhibits significantly enriched oxidative muscle fibers and dramatically decreased glycolytic muscle fibers (Fig S3F-H) (Gan, Rumsey et al., 2013). Consistent with the remarkably reduced glycolytic metabolism seen in *PPARβ* TG mice, the percentage of Pax7^Hi^ SCs was significantly reduced in the TA muscle fibers of *PPARβ* TG mice compared to those of wild-type (WT) littermates (Fig 3I-K). In agreement with this finding, the mRNA levels of *Pax7*, *CXCR4* and *CD34* were also lower, but the levels of *MyoD* and *MyoG* were higher in Pax7 SCs freshly isolated from the TA muscle of *PPARβ* TG mice compared to those of WT mice (Fig EV3I). Moreover, the activation of Pax7 SCs isolated from the TA muscle fibers of both WT and *PPARβ* TG mice was comparatively analyzed by immunostaining the Pax7 SCs cultured for 18 hr in growth medium with MyoD antibody. Activation of Pax7 SCs isolated from the TA muscle fibers of *PPARβ* TG mice was significantly faster than that observed in WT littermates (Fig EV3J and K). Together, these results support the notion that glycolytic metabolism favors the establishment and maintenance of Pax7^Hi^ cells in mice.

### Muscle-released G-CSF is a metabolic niche factor required for establishment and maintenance of Pax7^Hi^ SCs in mice

Based on the above observations, we hypothesized that cytokines highly expressed and secreted by glycolytic muscle fibers might function as niche factors required to maintain Pax7^Hi^ SC subpopulation in aged mice. To identify the muscle-secreted factors responsible for the loss of Pax7 SCs during aging, we analyzed the global gene expression changes occurring within TA muscle during physiological aging. We found that *Csf3* gene encoding granulocyte colony-stimulating factor (G-CSF) was particularly interesting because it was significant downregulated in aged TA muscle than young TA muscle (Fig 4A). Also, *Csf3* was more highly expressed by glycolytic TA muscle versus oxidative Sol muscle in adult mice (Fig 4B), and was remarkably reduced in the TA muscle of *PPARβ* TG versus WT adult mice (Fig 4C), suggesting *Csf3* gene is predominantly expressed in glycolytic muscle fibers. To further corroborate this, we isolated single myofibers from Sol and glycolytic EDL muscle and directly examined *Csf3* gene expression in single myofibers by qRT-PCR (Fig 4D). The identity of each myofibers was verified by measuring *Myh1* and *Myh7* genes expression, respectively (Fig EV4A and B). Indeed, *Csf3* was highly expressed in glycolytic single myofibers isolated from EDL (Fig 4D). Together, we reasoned that G-CSF might be a Pax7 SCs niche factor required for the established enrichment of Pax7^Hi^ SCs in glycolytic TA muscle in adult mice.

**Fig 4.**
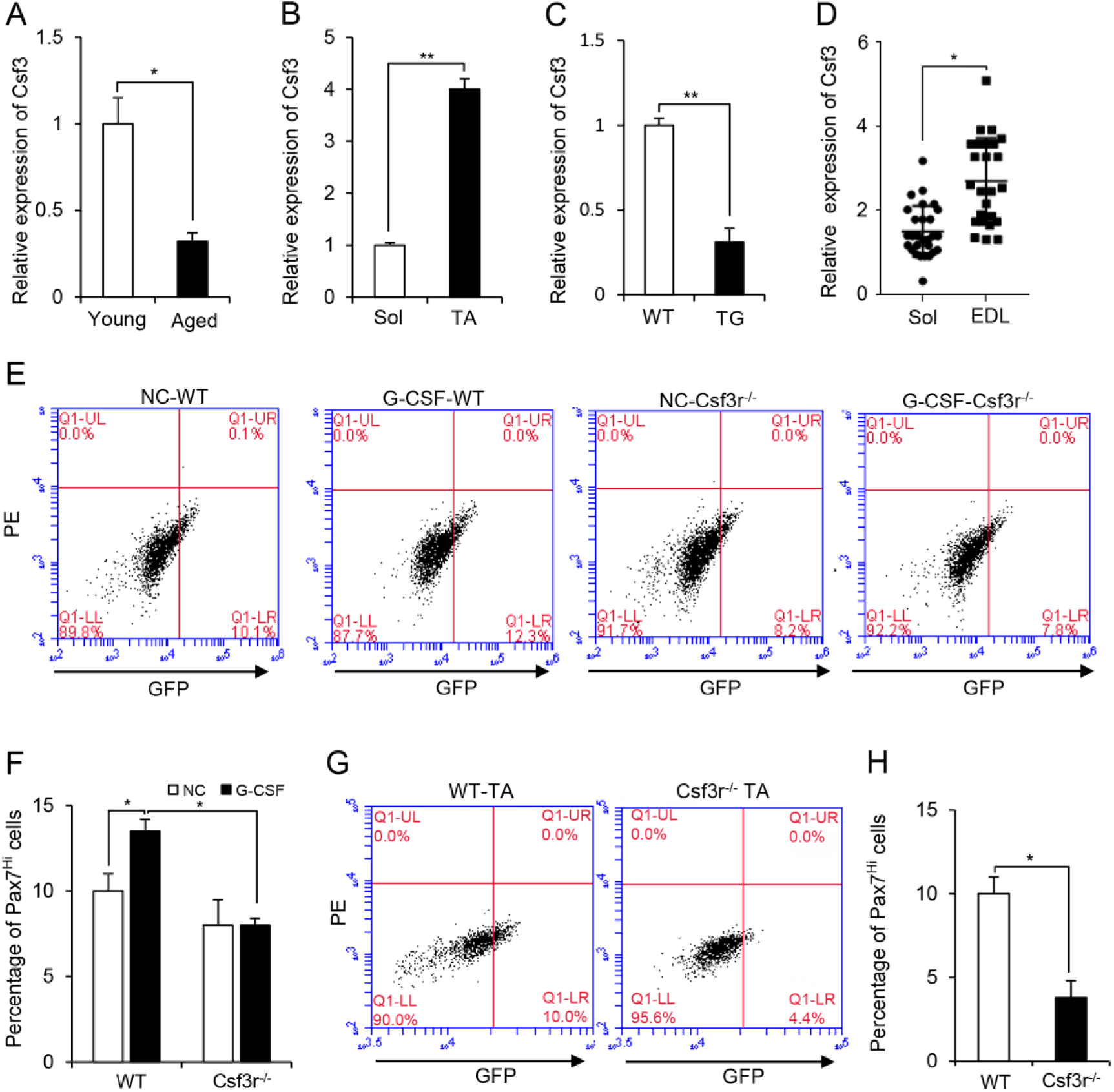
Muscle-released G-CSF is a Pax7 SC niche factor required for Pax7^Hi^ SCs. A Relative expression of *Csf3* gene in TA muscles from young (3-month-old) and aged (18-month-old) C57BL/6 mice, as determined by qRT-PCR. n = 5. *p< 0.05. Unpaired two-sided t-test. B Relative expression of *Csf3* gene in myofibers isolated from TA and Sol muscles were determined by qRT-PCR. *n* = 5. **p< 0.01. Unpaired two-sided t-test. C Relative expression of *Csf3* gene in TA muscles from *Pax7-nGFP;MCK-PPARβ-TG* and *Pax7-nGFP* WT mice, as determined by qRT-PCR. n = 5. **p< 0.01. Unpaired two-sided t-test. D Relative expression of *Csf3* gene in isolated single fiber from Sol muscle and EDL muscle were determined by qRT-PCR, each dot represent one single myofiber. n=25 in Sol group, n=23 in TA group. *p< 0.05. Unpaired two-sided t-test. E Representative FACS profile of Pax7 SCs treated with G-CSF in GM for 48 hr. Pax7 SCs were sorted from *Pax7-nGFP* WT and *Pax7-nGFP;Csf3r^-/-^*, respectively. PBS served as a negative control (NC). F Average percentages of Pax7^Hi^ SC subpopulations obtained from 3 independent FACS experiments, performed as described in panel (E). *p< 0.05. 2-way ANOVA. G Representative FACS profiles of Pax7 SCs from the TA muscles of 10-week-old *Pax7-nGFP;Csf3r^-/-^* or *Pax7-nGFP* WT mice. H The percentages of Pax7^Hi^ SCs in (G) were calculated. *n* = 3 for each genotype. *p< 0.05. Unpaired two-sided t-test.

To test this possibility, we freshly isolated Pax7 SCs from total muscle tissues of *Pax7-nGFP* mice, and treated the cells with either G-CSF or PBS (control) for 48 hr. G-CSF treatment significantly increased the percentage of Pax7^Hi^ SCs compared to the PBS control (Fig4E and F), indicating that G-CSF plays a functional role in modulating Pax7^Hi^ SCs *in vitro*. This was further supported by significant upregulation of Pax7 in SCs treated with G-CSF for 48 hr (Fig EV4C). To further confirm that G-CSF is a niche factor required for Pax7^Hi^ SCs establishment, we generated *Pax7-nGFP;Csf3r^-/-^*mice by crossing *Csf3r^-/-^*mice (G-CSF receptor KO mice) with *Pax7-nGFP* reporter mice. When Pax7 SCs isolated from the *Pax7-nGFP;Csf3r^-/-^*mice were treated with either G-CSF or PBS for 48 hr, G-CSF failed to increase the percentage of Pax7^Hi^ SCs from *Pax7-nGFP;Csf3r^-/-^*mice (Fig 4E and F). Consistent with these *in vitro* findings, when we compared the distribution of Pax7^Hi^ SCs in glycolytic TA of WT mice and *Pax7-nGFP;Csf3r^-/-^* mice, the percentage of Pax7^Hi^ SCs was significantly reduced in those of Pax7-nGFP;*Csf3r^-/-^* mice (Fig 4G and H). We next examined the mRNA levels of *Csf3* and marker genes for the muscle fiber type (*Myh4*) and muscle fiber metabolism (*HK2* and *PFK1*) in the TA muscles of *Pax7-nGFP;Csf3r^-/-^* and *Pax7-nGFP* WT mice. Our results ruled out the possibility that the observed effects in the *Pax7-nGFP;Csf3r^-/-^* mice were due to a reduction in*Csf3* expression or an alteration of muscle fiber metabolism (Fig EV4D and E). Moreover, we did not find any difference in the *Csf3* mRNA levels of immune cells sorted from the TA or Sol muscle fibers of *Pax7-nGFP;Csf3r^-/-^*mice (Fig EV4F and G). Collectively, these findings demonstrate that muscle-released G-CSF and its receptor on Pax7 SCs are required to establish the Pax7^Hi^ SC subpopulation in mice.

### Expression of *Csf3* gene encoding G-CSF is metabolically regulated by MyoD in muscle cells

We then asked if G-CSF is indeed a metabolic niche factor secreted by muscle fibers, then the *Csf3* gene encoding G-CSF should be metabolically regulated in muscle cells. To test this, we examined *Csf3* gene expression in C2C12 myotubes with enhanced glycolytic activity by culturing them in pyruvate-free medium (Fig EV5A and B) (Chen, Freinkman et al., 2016). The expression of *Csf3* gene was significantly increased in myotubes that exhibited higher glycolytic activity (Fig 5A), indicating that the transcription of *Csf3* gene was indeed regulated by the enhancement of glycolytic metabolism in myotubes. To determine how expression of *Csf3* gene is metabolically controlled by glycolytic activity of myofiber, we analyzed the 2-kb upstream of the transcriptional start site of the *Csf3* promoter and six E-boxes were found in this region (Fig EV5C), suggesting that metabolically-mediated expression of *Csf3* gene might be regulated by myogenic regulatory factors such as MyoD in myotubes. Therefore, we examined MyoD expression in myotubes cultured with pyruvate-free medium and found that enhanced glycolytic activity in the myotubes cultured with pyruvate-free medium also significantly elevated MyoD expression (Fig 5A). In addition, we also observed that similar to *Csf3*, *MyoD* was also more prominently expressed in glycolytic TA muscle fibers compared to oxidative Sol muscle fibers in mice (Fig EV5D and E). MyoD and *Csf3* had a similar expression pattern, highly expressed in the glycolytic single fibers isolated from EDL than in the oxidative single fibers from Sol muscles (Fig EV5F). To directly determine if MyoD metabolically regulates *Csf3* expression *in vivo*, expression of *Csf3* was assessed in TA muscles of MyoD-knockout (MyoD-KO) mice and WT mice. Consistently, levels of *Csf3* mRNA were significantly reduced in the glycolytic TA muscles of MyoD-KO mice compared to WT mice (Fig 5B), indicating that MyoD was a transcriptional factor for regulating *Csf3* expression in muscle cells. To further confirm this possibility, we overexpressed MyoD in C2C12 muscle cells and found that MyoD overexpression remarkably augmented *Csf3* transcription (Fig EV5G). MyoD-mediated transcription of *Csf3* gene was further corroborated by using a luciferase reporter system driven by the E-box-containing 2-kb region upstream of the *Csf3* promoter. Luciferase reporter activity was assayed in both C3H-10T1/2 fibroblasts and C2C12 cells transiently transfected with MyoD in the presence of the reporter construct. Forced expression of MyoD in either fibroblasts or C2C12 cells significantly activated the luciferase reporter gene driven by the 2-kb *Csf3* promoter, compared with the negative control (Fig 5C and EV5H). In addition, we examined luciferase reporter gene activity in the TA muscles of MyoD-KO and WT mice. Consistent with the results of our *in vitro* assays, the TA muscles of MyoD-KO mice showed significantly less luciferase activity than those of WT mice (Fig 5D). Together, our *in vitro* and *in vivo* analyses reveal that MyoD regulates the *Csf3* gene transcription in muscle cells.

**Fig 5.**
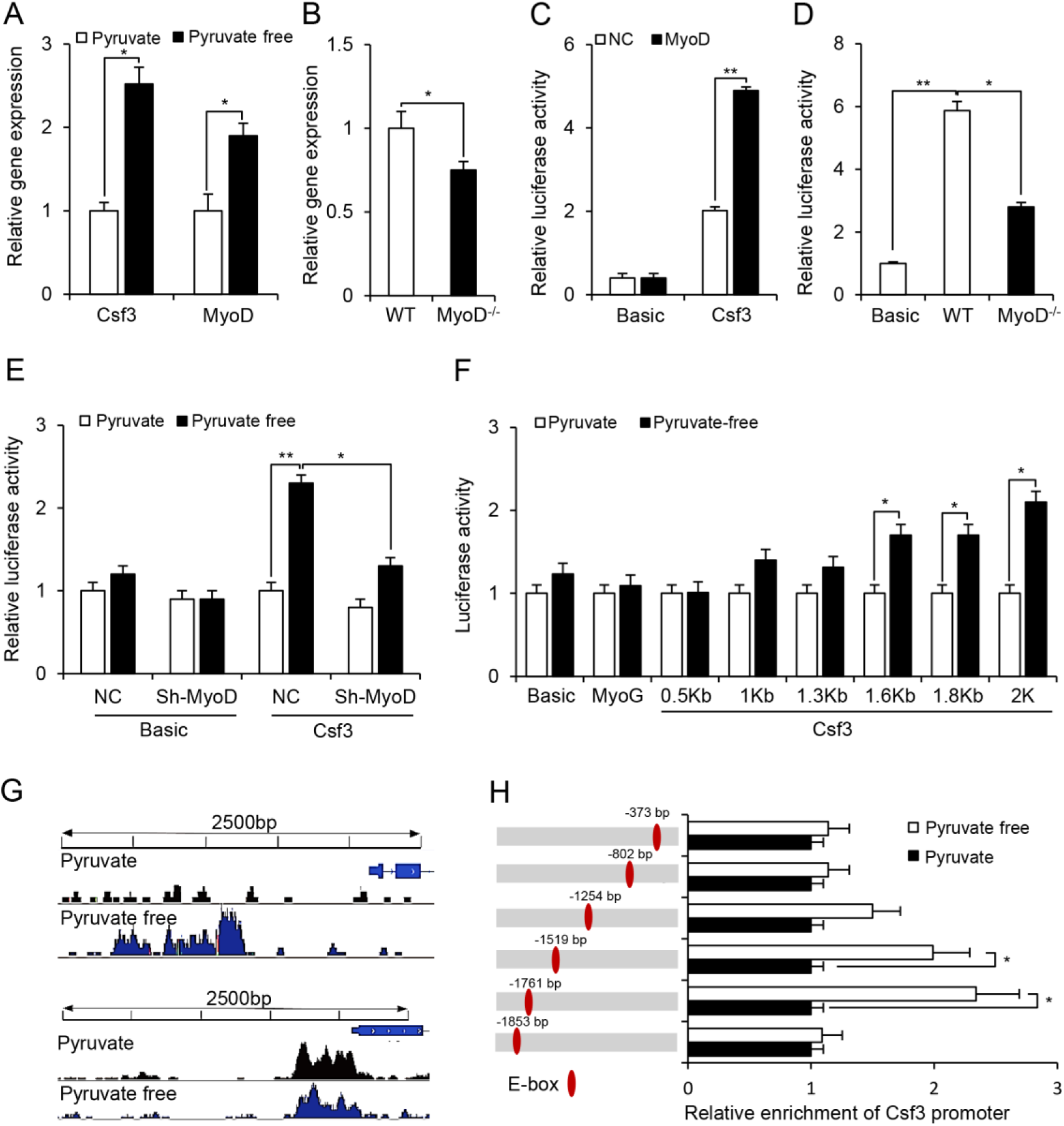
Expression of *Csf3*gene is metabolically regulated by MyoD in muscle cells. A Relative expression of *Csf3* and *MyoD* genes in C2C12 myotubes cultured in medium with or without pyruvate, as determined by qRT-PCR. Data were obtained from 3 independent experiments. *p< 0.05. Unpaired two-sided t-test. B Relative expression of *Csf3* gene in TA muscles from *MyoD^-/-^* mice and WT littermates were determined by qRT-PCR. *n* = 5 for each genotype. *p< 0.05. Unpaired two-sided t-test. C Relative *Csf3* promoter activities in C2C12 myotubes overexpressing MyoD were measured by dual luciferase assay. Empty vector served as a negative control (NC). Data were obtained from 3 independent experiments. **p< 0.01. 2-way ANOVA. D Relative *Csf3* promoter activities in the TA muscles of *MyoD^-/-^* and WT littermates were measured by dual luciferase assay. *n*=3 for each genotype. *p< 0.05, **p< 0.01. 1-way ANOVA. E Relative *Csf3* promoter activities in MyoD-knockdown C2C12 myotubes cultured in medium with or without pyruvate. Data were obtained from 3 independent experiments. *p< 0.05, **p< 0.01. 2-way ANOVA. F Relative activities of truncated *Csf3* promoters in C2C12 myotubes cultured in medium with or without pyruvate. Data were obtained from 3 independent experiments. *p< 0.05. Unpaired two-sided t-test. G ChIP-seq profiles of the *Csf3* and *MyoG* gene loci. Top to bottom: MyoD ChIP-seq profile in myotube cultured in presence of pyruvate (black signals) and absence of pyruvate (blue signals) on the promoter of *Csf3* (top panel) and *MyoG* (bottom panel). H ChIP assays were performed using chromatin from myotube cultured in medium with or without pyruvate. Chromatin was immunoprecipitated using antibodies against MyoD. The immunoprecipitated DNA was amplified using primers for *Csf3* gene promoter covering each indicated E-box, respectively. Data were obtained from 3 independent experiments.*p< 0.05. Unpaired two-sided t-test.

Next we examined whether the glycolytically-mediated transcription of the *Csf3* gene is regulated by MyoD in muscle cells. First, we checked *Csf3* expression in both MyoD-KD and control C2C12 myotubes with enhanced glycolytic activity. The metabolic reprogramming-induced upregulation of *Csf3* expression seen in control myotubes was completely abolished in the MyoD-KD myotubes (Fig EV5I and J), indicating that MyoD is required for the metabolically induced expression of *Csf3* gene in myotubes. To further confirm this observation, we assessed luciferase reporter gene activity driven by the *Csf3* promoter in MyoD-KD and control C2C12 myotubes cultured in the presence or absence of pyruvate. Indeed, significantly increased reporter gene activity was found to be induced by glycolytic metabolism only in control myotubes, but not in MyoD-KD myotubes (Fig 5E). Interestingly, enhanced glycolytic activity did not alter the expression of the endogenous *MyoG* gene (Fig EV5K) or the activity of a luciferase reporter gene driven by the *MyoG* proximal promoter (Fig EV5L), revealing that metabolically-mediated *Csf3* transcription is specifically regulated by MyoD in the muscle cells. Finally, functional analysis of *Csf3* gene promoter for identifying the E-boxes in its promoter required for MyoD-controlled metabolic transcription was performed by reporter gene assays with various truncated mutants of the *Csf3* gene promoter. Reporter assay showed that truncations of *Csf3* promoter containing the E-box (-1519bp, -1791bp, -1853bp) were response to glycolytic metabolism, suggesting that the E-boxes (-1519bp, -1791bp, -1853bp) were required for MyoD-mediated *Csf3* transcription (Figs 5F and EV5C). To further corroborate the observation, we performed MyoD ChIP-seq on differentiated myotubes cultured in the presence or absence of pyruvate. Significantly, we found that enhanced glycolytic metabolism enriched MyoD binding on *Csf3* promoter but not on *MyoG* promoter (Fig 5G). This was further confirmed by MyoD ChIP-PCR on *Csf3* promoter and MyoG promoter (Fig 5H and EV5M and N). Taken together, our results not only provide molecular evidence to confirm that G-CSF is a muscle fiber secreted niche factor but also most interestingly uncover an unexpected metabolic role for MyoD as a transcriptional factor in regulating *Csf3* gene expression in mature muscle.

### Muscle-derived G-CSF promotes the asymmetric division of Pax7 SCs

Next, we explored the molecular mechanism through which muscle-derived and MyoD-regulated G-CSF acts as a Pax7 SC niche factor to modulate the heterogeneity of Pax7 SCs. We first assessed the expression of *Pax7* in SCs following G-CSF treatment. Interestingly, at 24 hr post-treatment, there was no obvious change in *Pax7* mRNA levels in Pax7 SCs sorted from *Pax7-nGFP* mice (Fig EV6A), but these levels were significantly enhanced at 48 hr post-treatment (Fig EV4C). These results suggested that the G-CSF-mediated enrichment of the Pax7^Hi^ SC subpopulation most likely occurs through cell division of Pax7SCs rather than through an increase of *Pax7* expression in these cells.

As the self-renewal of the SCs was proposed to be regulated by asymmetric division (Kuang et al., 2007, Troy, Cadwallader et al., 2012) and G-CSF receptor was recently reported to be asymmetrically distributed in about 20% activated Pax7 SCs (Hayashiji, Yuasa et al., 2015), we hypothesized that G-CSF might mediate the heterogeneity of Pax7 SCs by promoting their asymmetric division via its interaction with the asymmetrically distributed G-CSFR on the Pax7 SCs. To test this possibility, we first performed time-lapse imaging of cell division in cultured Pax7 SCs sorted from *Pax7-nGFP* mice (Fig 6A-C and Movie1) and in single fibers isolated from the extensor digitorum longus (EDL) muscles of *Pax7-nGFP* mice (Fig EV6B-D and Movie 2). As expected, SCs underwent asymmetric division, each giving rise to one Pax7^Hi^ cell and one Pax7^Lo^ cell (Fig 6A-C and EV6B-D). Next, we used several approaches to experimentally test our notion that G-CSF regulated the percentage of the Pax7^Hi^ SC subpopulation by promoting their asymmetric division. Firstly, we assayed co-segregation of template DNA strands in the Pax7 cells. The TA muscles of 10-week-old *Pax7-nGFP* mice were injured by intramuscular injection of CTX. EdU labeling was used to monitor the co-segregation of template DNA (72 hr post injury), and BrdU was added to ensure that cells continued to divide during this period (8 hr post EdU) (Fig 6D). We ensured that all of the cells were EdU positive cells by performing immunostaining of EdU at T1 (Fig EV6E). To determine whether Pax7 cell displayed asymmetric division in response to G-CSF, we sorted Pax7 SCs and treated them with G-CSF in growth medium for 12 hr to complete cell division. In this paradigm, template DNA-retaining and -excluding cells would be EdU-positive or -negative, respectively (Fig 6E). We found that 12.5% of cells generated from Pax7 SCs were EdU-negative, suggesting that a subpopulation of Pax7SCs underwent co-segregation of template DNA (Fig 6F). Notably, the EdU-negative daughter cells were primarily Pax7^Lo^ SCs (Fig 6E). Significantly, G-CSF treatment increased the percentage of template DNA co-segregation in Pax7 SCs (Fig 6F), suggesting that G-CSF maintained Pax7^Hi^ SCs by promoting the asymmetric divisions.

**Fig 6.**
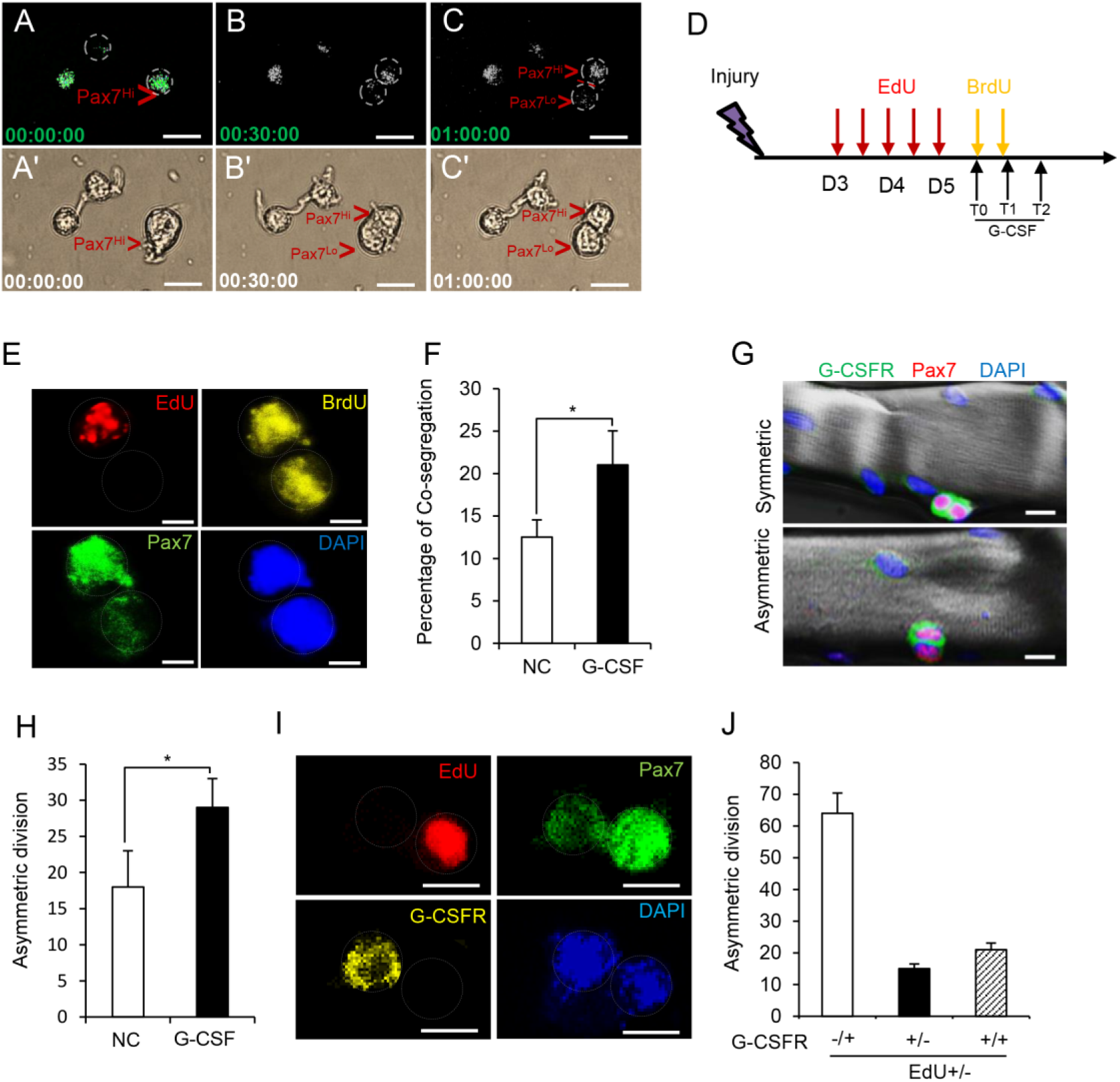
Muscle-derived G-CSF mediates the maintenance of the Pax7^Hi^ SC subpopulation by promoting the asymmetric division of Pax7 SCs. A-C Time-lapse imaging was used to trace the division of Pax7 SCs sorted from Pax7-nGFP mice. A-C were filmed in GFP channels and A’-C’ were filmed in bright channels, n=4. Scale bar represents 20 µm. D The timing of EdU and BrdU injections (8 hr apart) used to define TDSS. E Representative views showing the co-segregation of template DNA strands during the cell division of FACS-resolved Pax7 SCs from *Pax7-nGFP* mice pulse-labeled with EdU and BrdU, as characterized by EdU and BrdU staining. When old template DNA strands co-segregate, one daughter cell (EdU+/BrdU+) is a Pax7^Hi^ SC and the other (EdU-/BrdU+) is a Pax7^Lo^ SC. DAPI (blue) was used to visualize nuclei. Scale bar represents 5 µm. F The percentages of EdU+/BrdU+ and EdU-/BrdU+ daughter cells observed during the cell divisions of Pax7 SCs treated with G-CSF for 24 hr. PBS treatment served as a negative control (NC). The Pax7 SCs were FACS-resolved from *Pax7-nGFP* mice pulse-labeled with EdU and BrdU. Data were obtained from 3 independent experiments. *p< 0.05. Unpaired two-sided t-test. G Representative views showing the separation of G-CSFR signals (green) during the cell division of Pax7 SCs on *ex vivo*-cultured single muscle fibers. DAPI (blue) indicates nuclei. Scale bar represents 10 µm. H The percentages of Pax7 SCs that show asymmetric separation of G-CSFR during cell division on *ex vivo*-cultured single fibers treated with G-CSF were calculated from 3 independent experiments. PBS was served as a negative control (NC).*p< 0.05. Unpaired two-sided t-test. I Representative views of the asymmetric division of G-CSFR (yellow) and co-segregation of template DNA strands (EdU-) during the cell division of FACS-resolved Pax7 SCs from *Pax7-nGFP* mice that had been pulse-labeled with EdU and cultured in GM for 24 hr. DAPI (blue) indicates nuclei. Scale bar represents 10 µm. J Correlation between the asymmetric division of G-CSFR in daughter cells and the co-segregation of EdU-labeled template DNA during cell division. EdU+/-GCSFR-/+ represents a doublet in which one daughter is G-CSFR-negative and EdU positive (Pax7^Hi^) while the other is G-CSFR-positive and EdU negative (Pax7^Lo^).

As G-CSFR was recently reported to be asymmetrically distributed in about 20% of activated Pax7 SCs (Hayashiji et al., 2015), we reasoned that the asymmetric distribution of G-CSFR response to G-CSF mediated asymmetric division. Firstly, single fibers isolated from the EDL muscles of *Pax7-nGFP* mice were treated with G-CSF for 48 hr and immunostained with anti-G-CSFR (Fig 6G). Notably, G-CSF treatment significantly increased the percentage of asymmetrically dividing Pax7 SCs, as characterized by the asymmetric distributions of G-CSFR (Fig 6H).To further substantiate this observation, we calculated the percentage of EdU^+/-^ doublets with asymmetric distribution of G-CSFR. Indeed, the EdU^+/-^ doublets exhibited a lower percentage of G-CSFR^+/+^ cells with symmetrically distributed G-CSFR, and a significantly higher percentage of G-CSFR^-/+^ cells with asymmetric distribution of G-CSFR (Fig 6I and J). The frequency of asymmetric G-CSFR distribution was consistent with the co-segregation of template DNA. Together, these results demonstrate that G-CSF promotes the asymmetric division of Pax7 SCs.

### The G-CSF replenishes Pax7^Hi^ cells by stimulating asymmetric division of Pax7^Mi^ cells

Since reduced Pax7^Hi^ cells in aged mice is correlated with fiber metabolism shift from glycolytic to oxidative, we then test whether enhanced glycolytic fiber metabolism could rejuvenate Pax7^Hi^ cells in aged mice. As endurance exercise can significantly increase the glycolytic activity of muscle fibers (Heath, Gavin et al., 1983), we examined percentage of Pax7^Hi^ SCs in TA muscles of aged mice in which glycolytic muscle metabolism was enhanced by endurance exercise. As expected, exercise significantly augmented the glycolytic activity in the TA muscles of aged mice compared to those of sedentary aged mice (Fig EV7A). Notably, FACS analysis revealed that the percentage of Pax7^Hi^ SC subpopulation was dramatically increased in the TA muscles of exercised aged mice compared to sedentary aged mice (Fig 7A and B). Most significantly, we found that the percentage of Pax7^Hi^ SCs in the TA muscles of exercised aged mice was replenished almost to the level seen in untrained young mice (Fig 7A and B). These results indicate that enhanced glycolytic metabolism of myofiber rejuvenate Pax7^Hi^ cells in aged mice.

**Fig 7.**
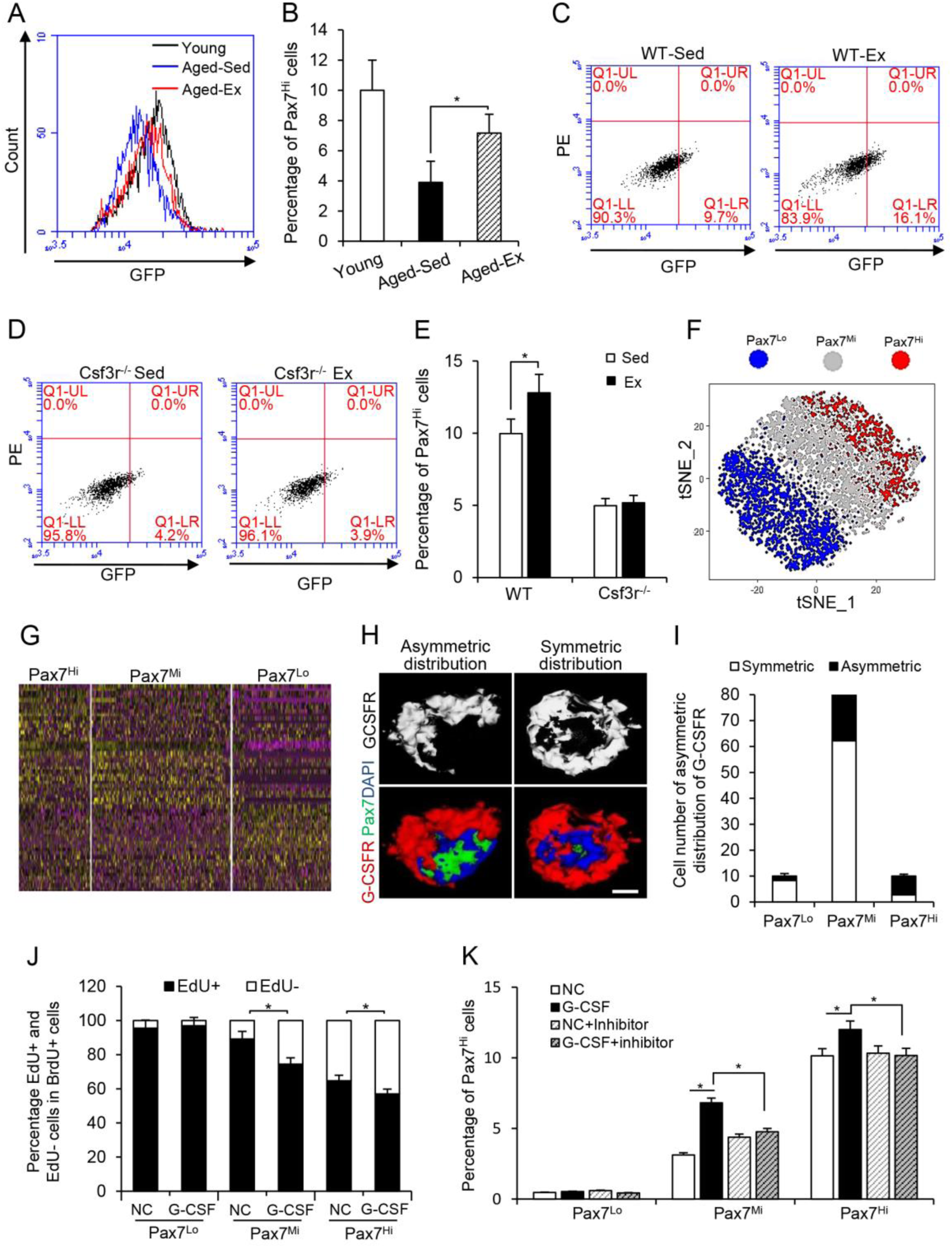
G-CSF replenishes Pax7^Hi^ cells by stimulating asymmetric division of Pax7^Mi^ cells. A Representative FACS profile of Pax7 SCs sorted from the TA muscles of young, aged exercised (aged-ex), and aged sedentary (aged-Sed) of Pax7-nGFP mice. B The percentages of Pax7^Hi^ SCs in A were calculated. *n* = 5 for each group. *p< 0.05. 1-way ANOVA. C-D Representative FACS profiles of Pax7 SCs from the TA muscles of *Pax7-nGFP;Csf3r^-/-^* or *Pax7-nGFP* WT mice subjected to exercise (Ex). Sedentary (Sed) mice served as a control. *n* = 5. E The percentages of Pax7^Hi^ SCs in C and D were calculated. *n* = 3. *p< 0.05. Unpaired two-sided t-test. F Unsupervised clustering of Pax7^Hi^, Pax7^Mi^ and Pax7^Lo^ cells visualized with tSNE. Each point was a single cell colored by cluster assignment. G Heatmaps of normalized signal show Pax7^Hi^, Pax7^Mi^, Pax7^Lo^ by top genes (columns) for individual cells (rows). H Representative views of the asymmetric and symmetric distributions of G-CSFR (red) in FACS-resolved Pax7 SCs obtained from *Pax7-nGFP* mice and cultured in GM for 24 hr. DAPI (blue) indicates nuclei. Scale bar represents 2.5 µm. I The percentages of Pax7 SCs with asymmetric distribution of G-CSFR among FACS-resolved Pax7^Hi^, Pax7^Mi^, and Pax7^Lo^ SCs cultured in GM for 24 hr. J The percentages of EdU+/BrdU+ and EdU-/BrdU+ daughter cells observed during the cell divisions of Pax7^Hi^, Pax7^Mi^, and Pax7^Lo^ SCs treated with G-CSF for 24 hr. PBS served as a negative control (NC). The three subpopulations of Pax7 SCs were FACS-resolved from *Pax7-nGFP* mice pulse-labeled with EdU and BrdU. Data were obtained from 3 independent experiments. *p< 0.05. Unpaired two-sided t-test. K The percentages of Pax7^Hi^ SCs after the three subpopulations of Pax7 SCs were treated with G-CSF for 24 hr in the presence or absence of a Stat3 inhibitor, as calculated from 3 independent experiments. *p< 0.05. 2-way ANOVA.

Given that exercise replenished Pax7^Hi^ cells through enhanced glycolytic metabolism of myofiber in aged mice (Fig 7A and B) and more interestingly, we also found that expression of *Csf3* was significantly reduced in TA muscles of aged mice and exercise significantly restored *Csf3* expression in the same fibers of aged mice (Fig EV7B). Based on those observations, it is conceivable that exercise-induced G-CSF might functionally restore Pax7^Hi^ cells in TA muscles of aged mice. To test this, we subjected *Pax7-nGFP;Csf3r^-/-^*and *Pax7-nGFP* WT mice to endurance exercise. Indeed, exercise significantly augmented the *Csf3* RNA levels of TA muscles from both exercised *Pax7-nGFP* WT and *Pax7-nGFP;Csf3r^-/-^* mice compared to those of sedentary *Pax7-nGFP* WT and *Pax7-nGFP;Csf3r^-/-^* mice (Fig EV7C). However, the percentage of Pax7^Hi^ SCs was only increased in the TA muscles of exercised *Pax7-nGFP* WT mice, but not in those of exercised *Pax7-nGFP;Csf3r^-/-^* mice (Fig7 C-E). Collectively, our results reveal that muscle-derived G-CSF acts as a metabolic niche factor required for maintaining the Pax7^Hi^ SC subpopulation in physiologically aged mice. These results indicate that enhanced glycolytic metabolism of myofiber rejuvenate Pax7^Hi^ cells in aged SCs by regulating *Csf3* expression.

Next, we asked from which subpopulations Pax7 SCs (Pax7^Lo^, Pax7^Mi^ and Pax7^Hi^) contribute to the replenishment of Pax7^Hi^ SCs in aged mice after exercise. For this purpose, we further sorted Pax7^Mi^ from *Pax7-nGFP* mice, and performed single cell RNA-seq. We profiled 5212 Pax7^Mi^ cells and the detectable genes ranged approximately from 1000 to 2000 in individual cells. Then we combined the data of Pax7^Hi^ and Pax7^Lo^ (Fig 1A) with the data of Pax7^Mi^ and visualized the cells in two dimensions according to their expression profiles by t-SNE projections. Our result showed that Pax7^Mi^ cells are more closed to Pax7^Hi^ cells (Fig 7F). Heatmaps of SCs profiles revealed normalized expression of the top variable genes in each subtype, and the expression pattern of Pax7^Mi^ cells were similar with Pax7^Hi^ cells (Fig 7G). Also, the markers of Pax7^Hi^ cells in Pax7^Mi^ cells were higher than Pax7^Lo^ cells (Fig EV7D). Hence, we reasoned that both Pax7^Hi^ and Pax7^Mi^ cells response to G-CSF to undergo asymmetric division. Consistently, we found that Pax7^Mi^ SCs had more cells with asymmetric distribution of G-CSFR, followed by the Pax7^Hi^ subpopulation, and then the Pax7^Lo^ cell subpopulation, which had a very low percentage (Fig 7H and I).To further determine which cell subpopulation(s) displayed asymmetric division in response to G-CSF, we used FACS to sort Pax7^Hi^, Pax7^Mi^ and Pax7^Lo^ SCs and treated them with G-CSF, respectively. We found that 36% of cells generated from Pax7^Hi^ SCs and 10% of those generated from Pax7^Mi^ SCs were EdU-negative, suggesting that most Pax7^Hi^ SCs and some Pax7^Mi^ SCs underwent co-segregation of template DNA (Fig 7J), indicating G-CSF treatment increased the percentage of template DNA co-segregation in both Pax7^Hi^ and Pax7^Mi^ SCs (Fig 7J). To further confirm these observations, we used flow cytometry to analyze template DNA co-segregation. After the cells finished their first cell division *in vitro*, ∼56% of the Pax7^Hi^ daughter cells were EdU^+^/BrdU^+^ and ∼44% were EdU^-^/BrdU^+^, whereas ∼18% of the Pax7^Mi^ generated cells were EdU^-^/BrdU^+^ (Fig EV7E). We also found that G-CSF treatment increased the percentage of template DNA co-segregation in both Pax7^Hi^ and Pax7^Mi^ SCs (Fig EV7E). As Pax7^Hi^ SCs generate distinct daughter cell fates by asymmetrically segregating template DNA strands to the stem cell (Rocheteau et al., 2012), only the Pax7^Hi^ cells themselves are not sufficient to enrich Pax7^Hi^ cells. The remarkable number of Pax7^Mi^ SCs with asymmetric distribution of G-CSFR provides a molecular basis for the G-CSF-mediated enrichment of Pax7^Hi^ SCs, which are generated through the asymmetric division of Pax7 SCs from the Pax7^Mi^ SC subpopulation. Finally, we examined the signaling pathway(s) involved in the G-CSF-mediated enrichment and maintenance of Pax7^Hi^ SCs. As G-CSF is known to activate the Stat3 signaling pathway in Pax7 SCs (Hara, Yuasa et al., 2011), we tested whether G-CSF regulated Pax7^Hi^ subpopulation through this pathway.

As reported, G-CSF activated Stat3 pathway and upregulated the downstream target genes in Pax7 SCs, but the Stat3 inhibitor significantly blocked effects of G-CSF (Fig EV7F and G). FACS-sorted Pax7^Hi^, Pax7^Mi^, and Pax7^Lo^ SCs were further treated with G-CSF in the presence or absence of the Stat3 inhibitor 5,15 DPP. G-CSF enriched the Pax7^Hi^ cell subpopulations in both Pax7^Hi^ and Pax7^Mi^ cell cultures, as indicated above, but treatment with the Stat3 inhibitor significantly abolished this enrichment (Fig 7K). These data indicate that G-CSF enriches Pax7^Hi^ cells through the G-CSF-G-CSFR-Stat3 axis. Collectively, our results offer multiple lines of experimental evidence showing that the G-CSF/G-CSFR/Stat3 axis is indispensably required to establish Pax7^Hi^ SC subpopulation in mice, and that it acts by promoting the asymmetric division of Pax7SCs.

## Discussion

Cell metabolism has been shown to intrinsically and cell-autonomously regulate cellular functions in various types of cells, especially in cancer cells (Carey, Finley et al., 2015, Moussaieff, Rouleau et al., 2015, Ryall, Dell’Orso et al., 2015). However, it was not previously known whether tissue metabolism plays an extrinsic and non-cell-autonomous role in modulating cell functions *in vivo*. A recent *in vitro* study reported that stem cell functions are modulated by the metabolic interplay between supporting Paneth cells and intestinal Lgr5^+^ crypt base columnar cell (Lgr5+CBCs) (Rodriguez-Colman, Schewe et al., 2017, Roper & Yilmaz, 2017). Here, we used various mouse genetic models to show for the first time that muscle fiber metabolism plays an *in situ* metabolic niche role in establishing and maintaining Pax7 SC heterogeneity in adult and physiologically aged mice. Thus, we reveal that the local metabolic activity of a tissue can provide *in situ* niche signaling to regulate stem cell functions *in vivo*. To our knowledge, our findings provide the first evidence that a tissue metabolism per se can act as a metabolic niche in regulating behaviors of stem cells during development and aging in a living organism.

Skeletal muscle is the most abundant endocrine organ and exerts its functional roles by secreting various factors (Pedersen & Febbraio, 2008, Pedersen & Febbraio, 2012). Muscle SCs are located between the sarcolemma and the basal lamina of the muscle fibers, which provide an immediate niche for the SCs by secreting different kinds of factors (Chakkalakal, Jones et al., 2012). However, only a few of the niche factors have been identified. For example, FGF2 was an aged muscle fiber-released cytokine that acts locally as an extrinsic factor to regulate muscle stem cell quiescence in aged mice (Chakkalakal et al., 2012). Actually, the muscle SCs are directly associated with two types of metabolically different fibers: glycolytic fibers and oxidative fibers. This locally metabolic environment of muscle fibers with different metabolic activity has been considered as a metabolic stem cell niche; however, this metabolic niche hypothesis has not been investigated experimentally. Using this unique metabolic system, we herein report on identification and molecular characterization of the metabolic niche factor G-CSF. We show that the G-CSF is highly secreted from glycolytic muscle fibers and its expression is metabolically regulated by MyoD in muscle fibers. Functionally, the muscle fiber-secreted G-CSF is required for establishing and maintaining the Pax7^Hi^ SC subpopulation in adult and physiological aged mice. Mechanistically, the muscle fiber-released G-CSF promotes the asymmetric division of Pax7^Hi^ and Pax7^Mi^ SCs by interacting with its receptor, G-CSFR, on Pax7 SCs in mice. To our knowledge, this is the first identified metabolic niche factor which is functionally required for regulating stem cell heterogeneity. The significance of our findings in general is that we provide molecular mechanism to conceptually prove metabolic niche hypothesis.

An unexpected finding of this study is the transcriptional activity of MyoD in mature muscle in mice. MyoD has long been regarded only as a master transcription factor with critical roles in controlling myogenic lineage specification during embryonic skeletal muscle development and activation of Pax7 SCs in response to muscle injury in adult mice (Cornelison, Olwin et al., 2000, Megeney, Kablar et al., 1996). Herein, we intriguingly found that MyoD predominately expressed in glycolytic muscle and metabolically regulated transcription of muscle-secreted G-CSF gene *Csf3* in mature muscle. Mechanistically, we show that enhanced glycolytic metabolism of myotube significantly enriched MyoD binding on *Csf3* promoter but not on the promoter of myogenic differentiation gene *MyoG*. These results for the first time reveal that MyoD is a multifunctional transcription factor involved in regulating expression of either myogenic genes or metabolic-regulated genes in mature muscle. A major question for future studies is how the specificity of MyoD transcriptional activity is achieved in different biological contexts. Better understanding of this question will be greatly facilitated by identification of MyoD-interacting cofactors in various biological settings. Taken together, these findings provide a framework to investigate the unanticipated and novel role of MyoD and examine the broad function of this cell-lineage specific transcription factor.

Heterogeneity is one hallmark of adult stem cells. However, it remains unclear how this heterogeneity is established and maintained during development and aging. In this report, using single cell RNA seq, we are the first to demonstrate that Pax7^Hi^ and Pax7^Lo^ muscle stem cells sorted based on the levels of Pax7 expression represent two distinct bona fide subpopulations in mice and Pax7^Mi^ cells were more similar to Pax7^Hi^ cells. Most strikingly, our approaches in this study allow us to reveal the dramatically decreased percentage of Pax7^Hi^ SCs in the glycolytic muscle fibers of physiologically aged mice (from 10% in adult mice to 2.7% in aged mice). Aging causes a deterioration of muscle function and regeneration that most likely reflects a decline in stem cell number and function. Pax7^Hi^ SCs are characterized as quiescent SCs with a high regenerative capacity, so the age-related reduction of Pax7^Hi^ SCs could account for the decline in muscle regeneration and repair in the aged mice. More remarkably, the reduction of the Pax7^Hi^ SC subpopulation in the muscle fibers of aged mice can be rescued by the exercise-induced up-regulation of G-CSF. A recent study showed that G-CSFR is asymmetrically distributed in about 20% of activated Pax7 SCs (Hayashiji et al., 2015). Interestingly, we found that the percentage of Pax7 SCs with asymmetrically distributed G-CSFR differed significantly among the three Pax7 SC subpopulations; it was highest in Pax7^Mi^, followed by Pax7^Hi^, and then Pax7^Lo^. In addition, we also found that Pax7^Mi^ cells are very similar to Pax7^Hi^ cells based on the gene expression signatures from single cell RNA sequencing. It therefore seems logical to propose a model in which Pax7^Mi^ SCs might represent an intermediate population of transitionally amplified Pax7 SCs that function as a reserve of Pax7 SCs from which active SCs are replenished, protecting the muscle stem cells from becoming exhausted under homeostasis and particularly following injury or during aging. Taken together our findings not only decipher a molecular mechanism that contributes to maintaining quiescent Pax7^Hi^ SCs in aged mice, but also suggest a subpopulation-based targeting strategy for treating age-related muscle loss (e.g., sarcopenia) or muscular dystrophy.

## Materials and Methods

### Mouse lines and animal care

*Pax7-nGFP* Tg mice were kindly gifted by Dr. Shahragim Tajbakhsh (Institute Pasteur, France). The Pax7-nGFP mice used throughout of this study were generated by crossing the C57BL/6J mice with the Pax7-nGFP Tg mice (C57BL6:SJL/J). *MCK-PPARβ* transgenic (TG) mice were kindly gifted by Dr. Zhenji Gan (Nanjing University, China). *Csf3r^-/-^* (#017838) and *MyoD^-/-^*(#002523) mice were obtained from the Jackson Laboratory. Mice were housed in an animal facility and given free access to water and standard rodent chow. All animal procedures were approved by the Animal Ethics Committee of Peking Union Medical College, Beijing (China).

### Fluorescence activated cell sorting (FACS)

Pax7 SCs from the skeletal muscles of *Pax7-nGFP*, *Pax7-nGFP;MCK-PPARβ* TG, and *Pax7-nGFP;Csf3r^-/-^*mice were fluorescently sorted as previously described (Wu et al., 2015). Briefly, mononuclear muscle-derived cells were isolated from the tibialis anterior (TA) and soleus (Sol) muscles of *Pax7-nGFP* reporter mice (3 month and 18 month) by dispase and collagenase digestion, filtered through 70-µm and 40 -µm cell strainers, and directly sorted with a BD Aria II Cell Sorting System. Three subpopulations of Pax7 SCs were sorted from Pax7-nGFP reporter mice by FACS based on intensity of GFP expression levels as previously described (Rocheteau et al., 2012). Briefly, Pax7^Hi^ and Pax7^Lo^ two subpopulations were sorted by FACS at opposite ends of the spectrum of GFP expression levels. They each corresponded to 10% of the total population and named as Pax7^Hi^ and Pax7^Lo^, respectively. Rest of 80% of SC in the middle were isolated and designated as Pax7^Mi^.

To sort immune cells from TA and Sol muscles, single-cell suspensions were prepared using dispase and collagenase, blocked with goat serum for 10 min, and co-incubated with CD11b-PECy7 (BD, 561098) and Ly-6G-FITC (BD, 561105) in DMEM supplemented with 2% FBS for 15 min at 4°C. The immunostained cells were briefly washed, passed through a 40-μm nylon mesh (Falcon), suspended at 10^3^-10^7^ cells/ml, and further separated with the BD Aria II. The sorting gates were strictly defined on the basis of mono-antibody-stained control cells and the forward- and side-scatter patterns obtained from the cells of interest in preliminary tests.

### Single cell RNA Seq using the 10x Genomics Chromium Platform

scRNA-seq libraries were prepared with the Single Cell 30 Reagent Kit as instruction from User Guide v2 (10x Genomics). Cellular suspensions of Pax7^Hi^, Pax7^Mi^ and Pax7^Lo^ were loaded on a Chromium Controller instrument (10x Genomics) to generate single-cell gel bead-in emulsions (GEMs), respectively. GEM-reverse transcriptions (GEM-RTs) were performed in a Veriti 96-well thermal cycler (Thermo Fisher Scientific). After RT, GEMs were harvested and the cDNAs were amplified and cleaned up with the SPRIselect Reagent Kit (Beckman Coulter). Indexed sequencing libraries were constructed using the Chromium Single-Cell 30 Library Kit (10x Genomics) for enzymatic fragmentation, end-repair, A-tailing, adaptor ligation, ligation cleanup, sample index PCR, and PCR cleanup. Sequencing libraries were loaded on a HiSeqX10 (Illumina). Reads were aligned to mm10 reference assembly. Primary assessment with this software for the Pax7^Hi^ sample reported 1469 cell-barcodes with 4466 median unique molecular identifiers (UMIs, transcripts) per cell and 1496 median genes per cell sequenced to 96.7% sequencing saturation with 313,083 mean reads per cell. Primary assessment with this software for the Pax7^Mi^ sample reported 5859 cell-barcodes with 3465 median unique molecular identifiers (UMIs, transcripts) per cell and 1260 median genes per cell sequenced to 94.9% sequencing saturation with 354,032 mean reads per cell. Primary assessment with this software for the Pax7^Lo^ sample reported 2982 cell-barcodes with 4317 median unique molecular identifiers (UMIs, transcripts) per cell and 1478 median genes per cell sequenced to 97.1% sequencing saturation with 300,302 mean reads per cell.

### Statistical method of Single cell RNA Seq

We used Cell Ranger version 1.3.1 (10x Genomics) to process raw sequencing data and Seurat suite version 2.0.0 for downstream analysis. The Seurat R package were used for graph-based clustering and visualizations, all functions mentioned were from this package or the standard R version 3.4.2 package unless otherwise noted and were used with the default parameters unless otherwise noted. Initially, we merged the three libraries by Seurat and we analyzed only cells (unique barcodes) that passed quality control processing (above) and expressed at least 500 genes and only genes that were expressed in at least 3 cells. We also removed cells with greater than 1% mitochondrial genes. We applied library-size normalization to each cell with NormalizeData. Normalized expression for gene i in cell j was calculated by taking the natural log of the UMI counts for gene i in cell j divided by the total UMI counts in cell j multiplied by 10,000 and added to 1. To reduce the influence of variability in the number of UMIs, mitochondrial gene expression between cells on the clustering, we used the ScaleData function to linearly regress out these sources of variation before scaling and centering the data for dimensionality reduction. Principle component analysis was run using RunPCA on the variable genes calculated with FindVariableGenes (x = (0.1,6), y = (0.5, 15) and then extended to the full dataset with ProjectPCA. Based on the PCElbowPlot result we decided to use 1 and 10 principle components (PCs) for the clustering of cells. We ran FindClusters to apply shared nearest neighbor (SNN) graph-based clustering to each sample (0.6).

### RNA seq

Total RNA was isolated from TA muscle of young (3-month-old) and aged (18-month-old) mice with Trizol reagent (Invitrogen). Sequencing libraries were generated using NEBNext super speed RNA Library Prep Kit for Illumina following the manufacturer’s recommendations. Raw-sequencing data were mapped to the mouse genome mm10 assembly using the HISAT with default parameters. DEGSeq45 was used to calculate the read coverage for each gene. Differentially expressed genes were filtered using a change greater than twofold and p-value (0.05) as a criterion for differential expression. Differentially expressed genes were validated using the iQ5 Multicolor Real-Time PCR Detection System (Bio-Rad). The primer sequences were designed using DNAMAN. Pax7-(s) CCGTGTTTCTCATGGTTGTG, (as)

GAGCACTCGGCTAATCGAAC; MyoD-(s) CAACGCCATCCGCTACAT, (as)

GGTCTGGGTTCCCTGTTCT; MyoG-(s)-CCATTCACATAAGGCTAACAC, (as)-

CCCTTCCCTGCCTGTTCC; Csf3-(s)-AGTGCACTATGGTCAGGACGAG, (as)

GGATCTTCCTCACTTGCTCCA; Nd1-(s) CATACCCCCGATTCCGCTAC, (as)

GTTTGAGGGGGAATGCTGGA; Co3-(s) ACCAATGATGGCGCGATGTA, (as)

GGCTGGAGTGGTAAAAGGCT; Co2-(s) CCGTCTGAACTATCCTGCCC, (as)

GAGGGATCGTTGACCTCGTC; Ptprb-(s) GCTGCCACGGCCCTT; (as)

CTCTGCCACTCCAGTCTGC; Pvalb-(s) ACACTGCAGCGCTGGTCATA, (as)

AGGAGTCTGCAGCAGCAAAGG; Rps28-(s) GGTGACGTGCTCACCCTATT, (as)

CCAGAACCCAGCTGCAAGAT.

### Chromatin Immunoprecipitation (ChIP)

ChIP analyses were performed on chromatin extracts from myotube cultured with or without pyruvate according to the manufacturer’s standard protocol (Millipore, Cat. #17-610) using antibodies against MyoD (Santa Cruz, SC-760). Briefly, cells were lysed in RIPA buffer (PBS, 1% NP-40, 0.5% sodium deoxycholate, 0.1% SDS) and centrifuged at 800 × g for 5 min. The chromatin fraction was sheared by sonication in 1.5 ml siliconized Eppendorf tubes. The resulting sheared chromatin samples were cleared for 1 hr, immunoprecipitated overnight, and washed in buffer I (20 mM Tris HCl [pH 8.0], 150 mM NaCl, 2 mM EDTA, 0.1% SDS, 1% Triton X-100), buffer II (20 mM TrisHCl [pH 8.0], 500 mM NaCl, 2 mM EDTA, 0.1% SDS, 1% Triton X-100), buffer III (10 mM TrisHCl [pH 8.0], 250 mM LiCl, 1% NP-40; 1% sodium deoxycholate, 1 mM EDTA), and Tris-EDTA (pH 8.0). All washes were performed at 4°C for 5 min. Finally, crosslinking was reversed in elution buffer (100 mM sodium bicarbonate [NaHCO3], 1% SDS) at 65℃ overnight. The resulting DNA were subjected to library construction for sequencing.

### Analysis of ChIP sequencing (ChIP-Seq)

ChIP-Seq data were obtained using an Illumina HiSeq 2000, with samples de-multiplexed via the Illumina pipeline, and mapped to the mouse genome (UCSC genome browser, mm9 version) using the Bowtie algorithm. ChIP-Seq data generated from mock DNA immunoprecipitates (input DNA) were used against the sample data in calling enriched regions and to control for the false-positive detection rate (FDR). MyoD peaks were called using MACS version 1.4.1, with p-value set to 10^-5^ for enrichment against the input genomic DNA, background reads were shuffled and randomly down sampled to adjust for the difference in read coverage between samples. Downstream analyses to generate intensity profile around the transcriptional start site (TSS), and correlative analysis with RNA-Seq data was completed using custom written codes in MATLAB. Fold enrichment was also quantified using qRT–PCR. The primer sequences were designed using DNAMAN. G-CSF Primer 1-(s) ATCACAAATGAAGGGCAGAG, (as) CAAGACTGCTTCTGTCTCTCC; G-CSF Primer 2-(s) ATGAGCAGAGATCGTCGGGA, (as) CACATTACCTCGATGTCGTG; G-CSF Primer 3-(s) TGTCCTCTCAAGCAGAGGCTAT, (as) GATGTTGAGGCATACCTGATG; G-CSF Primer 4-(s) CGCAAGATGTCTATCTG, (as) CCATGCCCGGCGAGATTTAATTC; G-CSF Primer 5-(s) CTTGTGCAGCTCATCAAGGC, (as) GTGGTGGGGATCTTTTGCTG; G-CSF Primer 6-(s) GCTACATTCTGAACGCTGCC, (as) GCCTTGATGAGCTGCACAAG.

### SDH and GPDH staining

For measurement of succinate dehydrogenase (SDH) activity, muscles were harvested and serial tissue cross-sections (10-µm) were cut at -20°C and adhered to glass coverslips. The coverslips were inverted and placed over a microscope slide reaction chamber. The tissue was first incubated in the dark at 23°C in a substrate -free blank solution consisting of 1 mM sodium azide, 1 mM l-methoxyphenazinemethosulfate (MPMS), 1.5 mM NBT, and 5 mM EDTA in 100 mM sodium phosphate buffer (pH 7.6). The reaction was allowed to proceed for 10 min to allow the nonspecific staining to plateau. The blank was then replaced with a substrate solution consisting of the above reagents plus 48 mM succinic acid. Images were captured every 3times for 10 min. For measurement of α-glycerophosphate dehydrogenase (α-GPDH) activity, serial sections (14-µm) were cut, adhered to glass coverslips, and distributed between two Coplin jars kept at -20°C. A blank solution consisting of 1 mM sodium azide, 1 mM MPMS, and 1.2 mM NBT in 100 mM sodium phosphate buffer (pH 7.4, 37°C) was added to one jar while a solution of the above reagents plus 9.3 mM α-glycerophosphate was introduced into the other for the substrate reaction. The tissue sections were incubated in the dark for 24 min at 37°C, the reactions were stopped by extensive rinsing with distilled water. The images were captured using a microscope (Olympus).

### Isolation and staining of single myofibers

Single myofibers were isolated from the EDL muscles of 3 month *Pax7-nGFP* mice by digestion with collagenase I (Sigma, C-0130), as previously described (Wu et al., 2015). Briefly, each muscle sample was incubated in 3 ml of 0.2% collagenase I in serum-free DMEM in a shaking water bath at 37°C for 45 -60 min. Digestion was considered complete when the muscle looked less defined and slightly swollen, with hair-like single fibers flowing away from the edges. The digested muscles were placed in a Petri dish, and myofibers were isolated under a microscope. Single fibers were placed in six-well plates pre-coated with horse serum, and then given 2 ml/well of fiber medium (DMEM supplemented with 20% FBS, 0.5% chick embryo extract, 10 pg/ml G-CSF, and penicillin-streptomycin). The fibers were cultured for 48hr at 37°C in a 5% CO _2_ atmosphere, fixed with 4% paraformaldehyde, and stained for G-CSFR. The fibers were then washed with PBS containing 0.1% BSA and incubated for 2 hr with fluorescein-conjugated secondary antibodies (Zhongshanjinqiao Corporation) and Hoechst or DAPI. For statistical analyses, the cells with symmetric and asymmetric distribution of G-CSFR were counted in at least 100 doublets per mouse. Five mice were assayed in each set of experiments.

### Immunofluorescence

FACS-resolved Pax7 SCs were seeded on collagen-coated glass slides in 24-well plates (2×10^4^ cells/cm^2^) in growth medium (F-10 containing 20% FBS) for 24 hr, fixed with 4% formaldehyde for 5 min, permeabilized in 0.1% Triton-X100 in PBS for 15 min at room temperature, and then blocked with 3% bovine serum albumin for 30 min. For BrdU immunostaining, the cells were unmasked with 2 N HCl for 20 min at room temperature and neutralized with 0.1 M sodium tetraborate. The cells were incubated with primary antibodies against MyoD (Santa Cruz, SC-760), G-CSFR (Santa Cruz, SC-9173), EdU (Invitrogen, C10640) and BrdU (Abcam, ab6326) overnight at 4°C. The cells were then washed with PBS containing 0.1% BSA and incubated for 2 hr with fluorescein-conjugated secondary antibodies (Zhongshanjinqiao Corporation) and Hoechst or DAPI. After several washes with PBS, the cells were examined under a fluorescence microscope (Olympus).

### Western blot analysis

TA and Sol muscles from C57BL/6 mice were homogenized in a buffer containing 50 mM Tris pH 7.5, 150 mM NaCl, 0.5% Nonidet P40, and protease and phosphatase inhibitors. The muscle homogenates were clarified by centrifugation at 12,000 × g for 10 min. Total proteins (40 µg) were resolved by SDS-PAGE, transferred to a polyvinylidene fluoride membrane, and immunoblotted with primary antibodies against MyoD (Santa Cruz, SC-760) and β-tubulin (Santa Cruz, SC-5274) overnight at 4°C. For Stat3 detection, the FACS -sorted Pax7 SCs were treated with G-CSF (10 pg/ml) in presence or absence of Stat3 inhibitor, 5,15 DPP (50 μM, Sigma-Aldrich), in growth medium for 48 hr. Subsequently, the nuclear and cytoplasmic fractions were further isolated from the treated SCs with the kit (Thermo, 78835). The nuclear (10 µg) and cytoplasmic protein (15 µg) were resolved by SDS -PAGE, transferred to a polyvinylidene fluoride membrane, and immunoblotted with primary antibodies against Stat3 (Abcam, ab19352), p-Stat3 (CST, 9154), histone H3 (Abcam, ab1791) and GAPDH (Millipore, Mab374). Membranes were washed for 30 min, incubated with horseradish peroxidase-conjugated secondary antibodies (Zhongshanjinqiao Corporation) for 1 hr at room temperature, and washed for 30 min. Each membrane was then placed into Detection Solution (Thermo), incubated for 1 min at room temperature, and exposed to X-ray film.

### RNA extraction and qRT-PCR

Total RNA was extracted from skeletal muscles using the TRIzol reagent (Invitrogen) and reverse transcribed with reverse transcriptase (Fermentas). Real-time quantitative PCR analyses were performed in triplicate using the Fast Eva Green qPCR Master Mix (BioRad). *GAPDH* was used as an internal control for qRT-PCR analyses.

### Treadmill

Young (3-month-old) and aged (18-month-old) *Pax7-nGFP;Csf3r^-/-^*mice, their wild-type littermates (*Pax7-nGFP*) were subjected to treadmill exercise using an Exer3/6 (Columbus Instruments). Mice were acclimated to treadmill running four times (every other day) before the test. Each mouse ran on the treadmill at 20°downhill, starting at a speed of 10 cm/s. After 3 min, the speed was increased by 2 cm/s to a final speed of 20 cm/s. Then the mice were allowed to run 25 min. After exercise training, the mice were sampled for purification and analysis of Pax7^Hi^ and Pax7^Lo^ SCs.

### Cell culture and treatments

FACS-resolved Pax7 SCs were cultured in F-10 medium containing 20% FBS and 2.5 ng/ml bFGF (Invitrogen) in the presence or absence of G-CSF (10 pg/ml, Santa Cruz) at 37°C in a 5% CO_2_ atmosphere. Fibroblasts (C3H-10T1/2, ATCC) were cultured in DMEM (Gibco) supplemented with 4.5 g/L glucose, 10% FBS, and 1% penicillin/streptomycin at 37°C in a 5% CO_2_ atmosphere. C2C12 cells (ATCC) were cultured in growth medium consisting of DMEM supplemented with 20% FBS and 1% penicillin/streptomycin. At 70–80% confluence, the C2C12 cells were switched to differentiation medium (DMEM with 2% horse serum). After 5 days of differentiation, the C2C12 myotubes were cultured in differentiation medium with or without pyruvate for 24 hr. For MyoD overexpression or knockdown, C2C12 myotubes were transiently transfected with pEGFPN1-MyoD or LV-sh-MyoD using Lipofectamine2000 (Invitrogen). The empty vector served as the negative control (NC).

### Luciferase reporter assay

To test promoter activity, a 2-kb sequence upstream of the *Csf3* gene was retrieved from the University of California Santa Cruz genome browser and cloned into the pGL3 Basic vector carrying the firefly luciferase gene (Promega). The generated pGL3-G-CSF-2k was used to transfect C3H-10T1/2 cells or C2C12 myotubes in differentiation medium with or without pyruvate. Empty pGL-3 vector was used as a negative control, and co-transfection with a Renilla luciferase plasmid (Promega) served as a transfection control. The results are expressed as the activity of firefly luciferase relative to that of Renilla luciferase. For promoter reporter gene assays *in vivo*, mouse TA muscles were injected with 20 μg of pGL3-G-CSF-2k plasmid DNA with electroporation. Briefly, plate electrodes were positioned on each side of the leg over the TA muscle, in contact with the skin. TA muscles were stimulated eight times (parameters: 100 V, duration 20 ms, interval 100 ms) with a stimulator (EM830, BTX). The same amount of empty pGL3 Basic vector served as a negative control. 5μg of Renilla luciferase plasmid were co-injected as a normalization control. Luciferase activity assays were performed with homogenates of TA muscle 24 hr after electroporation.

### Time-lapse imaging

Sorted Pax7 SCs or isolated single fibers from *Pax7-nGFP* mice were plated on a 24-well plate pre-coated with matrigel. After incubation overnight, the cells were filmed with a Real-Time Cell History Recorder (JuLi stage, NanoEnTek Inc., Korea) inside an incubator. Images were taken every 15 min using the bright and GFP channels. The raw data were transformed and were presented as a video.

### Template DNA strand segregation (TDSS)

Template DNA strand segregation was analyzed as previously reported (Rocheteau et al., 2012). Briefly, the TA muscles of *Pax7-nGFP* mice (3month) were injured by intramuscular injection of cardiotoxin (CTX; 50 μl of 10-μM CTX per TA muscle). Mice were injected intraperitoneally 3 days post injury with EdU (five times, 200 mg/injection, 8 hr apart) followed by injection of BrdU (twice, 8 hr apart). Then Pax7 SCs were sorted, cultured in DMEM for a further 24 hr, and immunostained with anti-BrdU (1:300, Abcam) and EdU (1:300, Invitrogen).

### Measurement of NAD

NAD^+^ and NADH were determined using commercially available kits (Biovision, Milpitas) according to the provider’s instructions.

### Statistical analysis

Data are presented as means +s.e.m. For statistical comparisons of two conditions, the two-tail Student’s *t*-test was used. Comparisons of multiple groups were made using a 1- or 2-way ANOVA. All experiments were repeated at least 3 times, and representative experiments are shown. Statistical analysis was performed in GraphPad Prism.

### Data availability

Related data were submitted to GEO with the accession number GSE134253.

## Acknowledgments

We greatly thank Yunning Geng and Huimin Liao from Bio-reach Co., Ltd., for their technical support of our time-pulse imaging experiments, and Dr. Shahragim Tajbakhsh and Dr. Zhenji Gan for providing the Pax7-nGFP mice and *PPARβ* transgenic mice. This work was supported by grants from the National Basic Research Program of China (2015CB943103, 2016YFA0100703), the National Natural Science Foundation of China (91540206, 31471289, 31471377), and the CAMS Initiative for Innovative Medicine (2016-I2M-1-017).

## Author contributions

DZ conceived the project, coordinated the study, and wrote the paper. HL designed and performed the experiments. QC performed single cell RNA library construction and ChIP DNA library construction. CL performed the screening of muscle-secreted factors. RZ and YXZ performed the luciferase reporter gene assays. QZ performed analysis of SCs from aged Pax7-nGFP mice. WT gave constructive discussion and helped to draft the manuscript. YZ helped draft the figure legends and experimental procedures of the manuscript. All authors approved the final version of the manuscript.

## Conflict of interest

The authors declare that they have no conflict of interest.

---

**Fig EV1.**
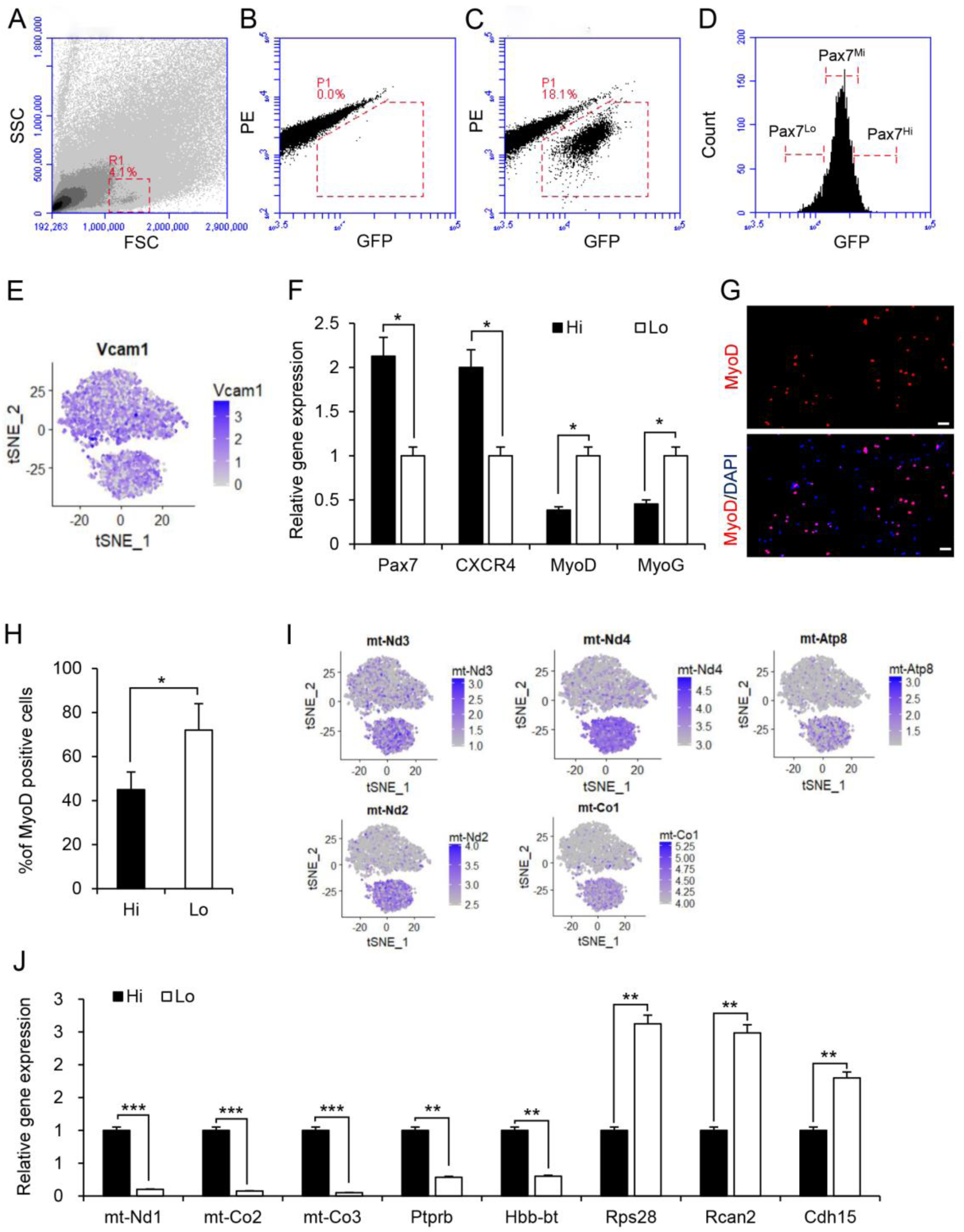
Characterization of FACS-sorted Pax7^Hi^ and Pax7^Lo^ cells. A-D Strategy used for FACS sorting GFP+ cells from*Pax7-nGFP* mice. Negative control for GFP gating was shown in B. Pax7^Hi^ and Pax7^Lo^ two subpopulations were sorted by FACS at opposite ends of the spectrum of GFP expression levels. They each corresponded to 10% of the total population and named as Pax7^Hi^ and Pax7^Lo^, respectively. Rest of 80% of SC in the middle were isolated and designated as Pax7^Mi^ (C and D). E Expression pattern of quiescent markers of satellite cells was visualized by t-SNE plots. F Relative expression of molecular markers for stemness and differentiation in FACS-resolved Pax7^Hi^ and Pax7^Lo^ SCs from the TA muscles of *Pax7-nGFP* mice, as determined by qRT-PCR. *n* = 5. *p< 0.05. Unpaired two-sided t-test. G Representative images of MyoD immunostaining (red) for FACS-resolved Pax7 SCs cultured in growth medium (GM) for 18 hr. DAPI was used to visualize nuclei (blue). Scale bar represents 20 µm. H The percentages of MyoD-positive cells in G were calculated from three independent experiments. *p< 0.05. Unpaired two-sided t-test. I Expression pattern of mitochondrial genes in Pax7^Hi^ and Pax7^Lo^ cells was visualized by t-SNE plots. J Relative expression of molecular markers in Pax7^Hi^ cells and Pax7^Lo^ cells FACS-resolved from *Pax7-nGFP* mice, as determined by qRT-PCR. *n* = 5. **p< 0.01. ***p< 0.001. Unpaired two-sided t-test.

**Fig EV2.**
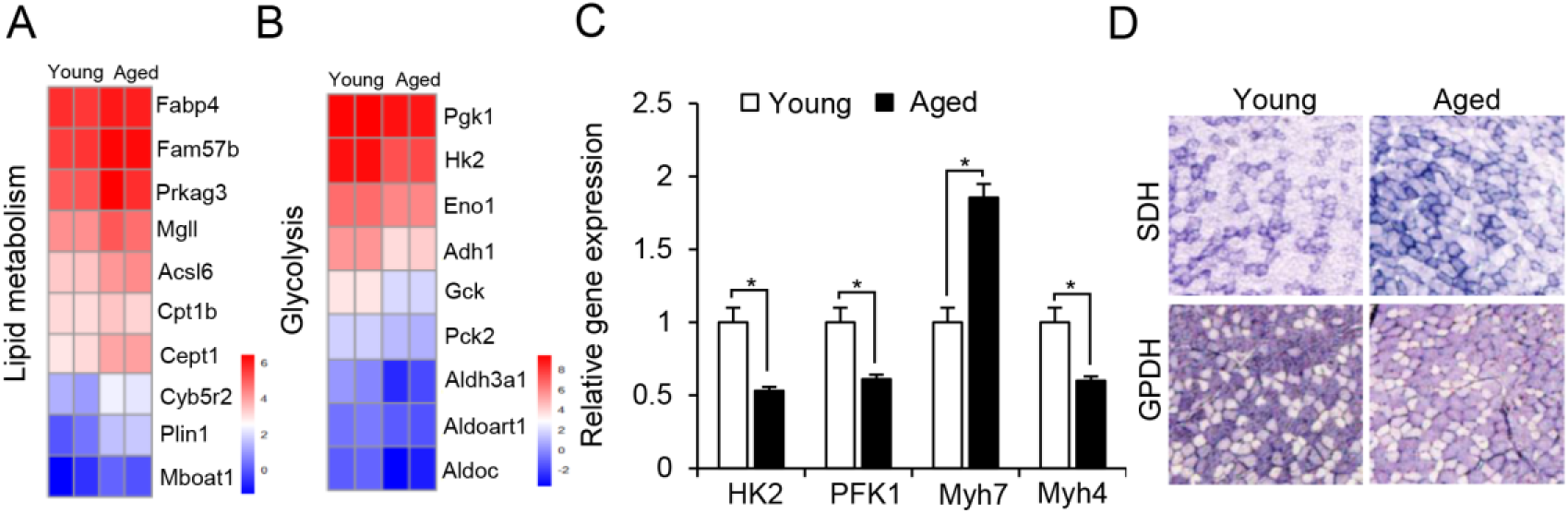
Muscle metabolism shift from glycolytic to oxidative activity during ageing. A,B Heatmaps indicated absolute gene expression (Log2[FPKM]) of specific metabolic regulators in aged and young muscle. Each gene listed had a mean fold change of greater than 1.5. C Relative expression of *PFK1*, *HK2, Myh7* and *Myh4* in TA muscles from young (3-month-old) and aged (18-month-old) mice, as determined by qRT-PCR. *n* = 5. *p < 0.05. Unpaired two-sided t-test. D Representative histochemical staining of α-GPDH and SDH enzymatic activity in TA muscle of aged and young mice.

**Fig EV3.**
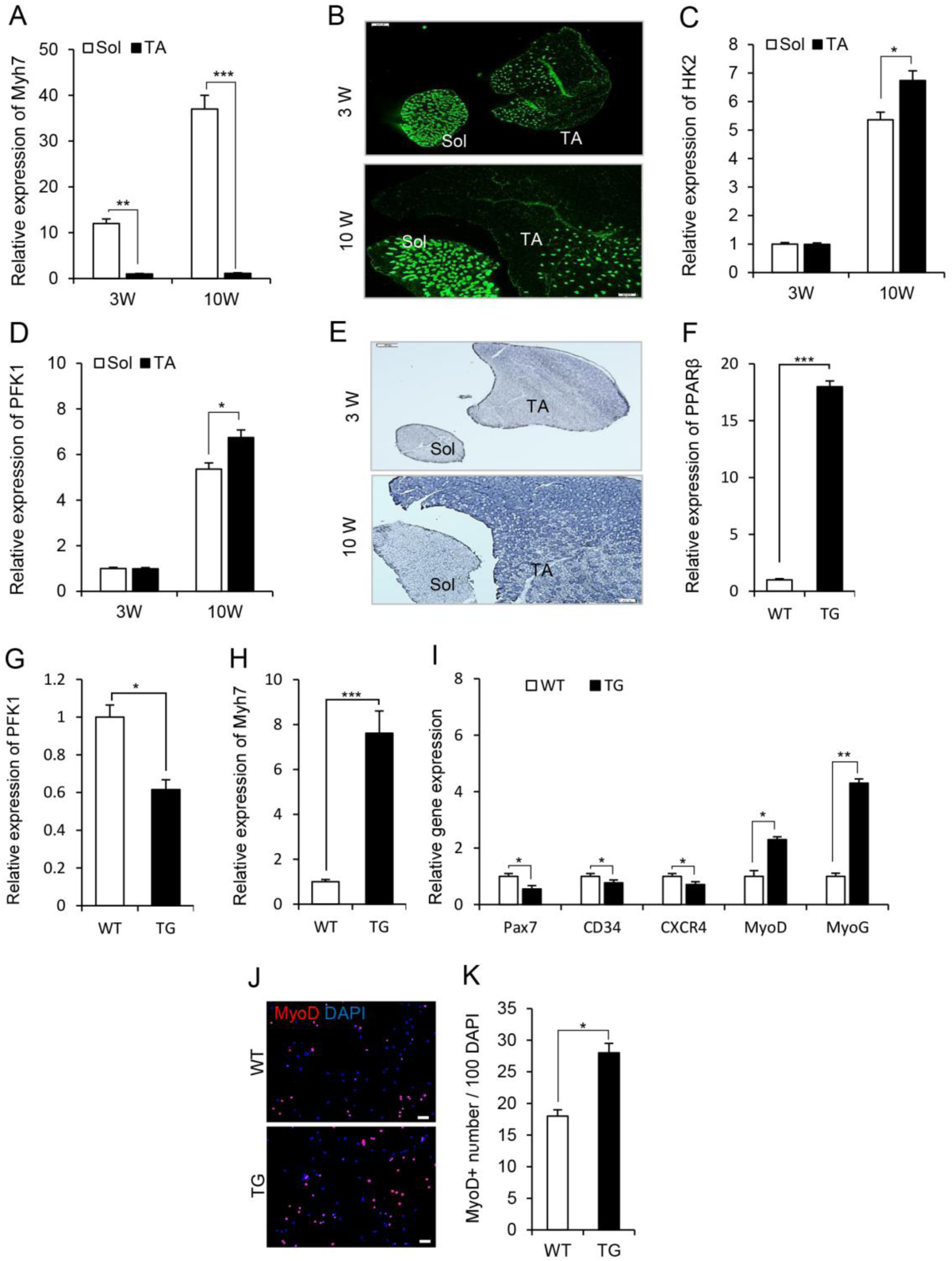
Fiber type composition and metabolic status in TA and Sol muscle and in TA muscle of *MCK-PPARβ-TG*. A Relative expression of *Myh7* in TA and Sol muscles from 3 weeks and 10 weeks C57BL/6 mice, as determined by qRT-PCR. *n* = 5. **p< 0.01. ***p < 0.001. Unpaired two-sided t-test. B Representative immunostaining of type I myofibers in TA and Sol muscle from 3 weeks and 10 weeks mice. C-D Relative expression of *HK2* (C) *and PFK1* (D) in TA and Sol muscles from 3 weeks and 10 weeks mice, as determined by qRT-PCR. *n* = 5. *p < 0.05. Unpaired two-sided t-test. E Representative histochemical staining of α-GPDH enzymatic activity in TA and Sol muscles from 3 weeks and 10 weeks mice. F-H Relative expressions of *PPARβ* (F), *PFK1* (G), and *Myh7* (H) in TA muscles from *Pax7-nGFP;MCK-PPARβ*-TG (TG) and *Pax7-nGFP* WT mice, as determined by qRT-PCR. *n* = 5. *p< 0.05. ***p < 0.001. Unpaired two-sided t-test. I Relative expression of molecular markers for stemness and differentiation in FACS-resolved Pax7 SCs from the TA muscle of *Pax7-nGFP;MCK-PPARβ* transgenic (TG) and *Pax7-nGFP* wild-type (WT) littermates, as determined by qRT-PCR. *n* = 5. *p< 0.05. **p < 0.01. Unpaired two-sided t-test. J Representative images of MyoD immunostaining (red) for FACS-resolved Pax7 SCs cultured in growth medium (GM) for 18 hr. DAPI was used to visualize nuclei (blue). Scale bar represents 20 µm. K The percentages of MyoD-positive cells in (J) were calculated from 3 independent experiments.*p < 0.05. Unpaired two-sided t-test.

**Fig EV4.**
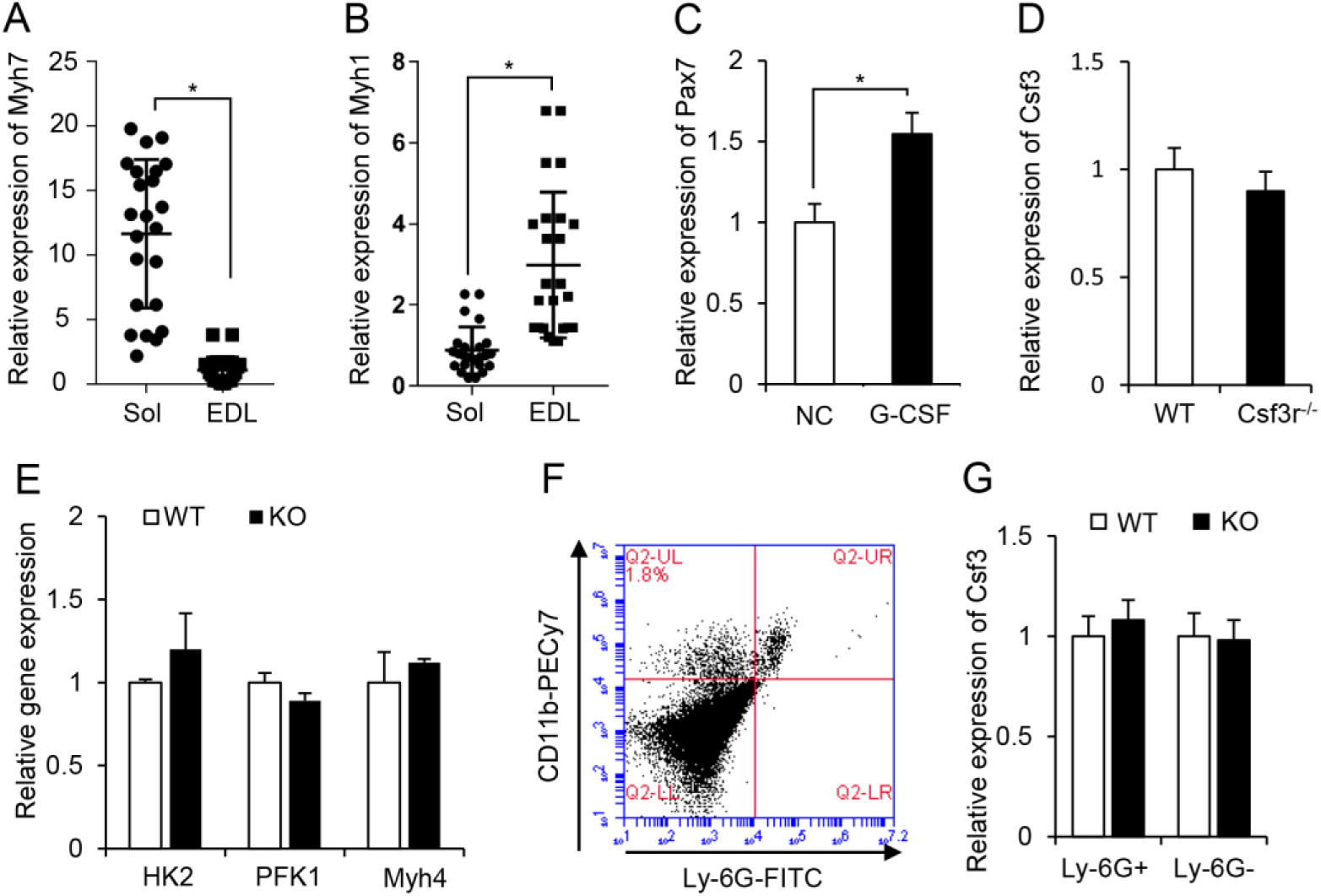
Muscle-released G-CSF is a Pax7 SC niche factor required for Pax7^Hi^ SCs. A Relative expression of *Myh7* gene in isolated single fiber from Sol and EDL muscles was determined by qRT-PCR, each dot represented one single myofiber. n=25 in Sol group. n=24 in EDL group.*p < 0.05. Unpaired two-sided t-test. B Relative expression of *Myh1* gene in isolated single fiber from Sol and EDL muscles was determined by qRT-PCR, each dot represented one single myofiber. n=25 in Sol group. n=24 in EDL group. *p < 0.05. Unpaired two-sided t-test. C Relative expression of *Pax7* in sorted Pax7 SCs treated with G-CSF in growth medium (GM) for 48 hr, as determined by qRT-PCR. Data were obtained from 3 independent experiments. *p< 0.05. Unpaired two-sided t-test. D Relative expression of *Csf3* gene in TA muscle from *Pax7-nGFP;Csf3r^-/-^* mice and *Pax7-nGFP* WT littermates, as determined by qRT-PCR. *n* = 5. E Relative expression of genes related to fiber type (*Myh 4*) and muscle metabolism (*HK2* and *PFK1*) in TA muscles from *Pax7-nGFP;Csf3r^-/-^* mice and *Pax7-nGFP* WT littermates, as determined by qRT-PCR. *n* = 4. F Representative FACS profiles of neutrophils and macrophages. Gating for CD11b+/Ly-6G+ (neutrophils) and CD11b+/Ly-6G-(macrophages) is indicated. FITC, GFP (488 channel); PEcy7, phycoerythrin (594 channel). G Relative expression of *Csf3* gene in the neutrophils and macrophages sorted from TA muscle of *Pax7-nGFP;Csf3r^-/-^* mice and *Pax7-nGFP* WT littermates, as determined by qRT-PCR. *n* = 4.

**Fig EV5.**
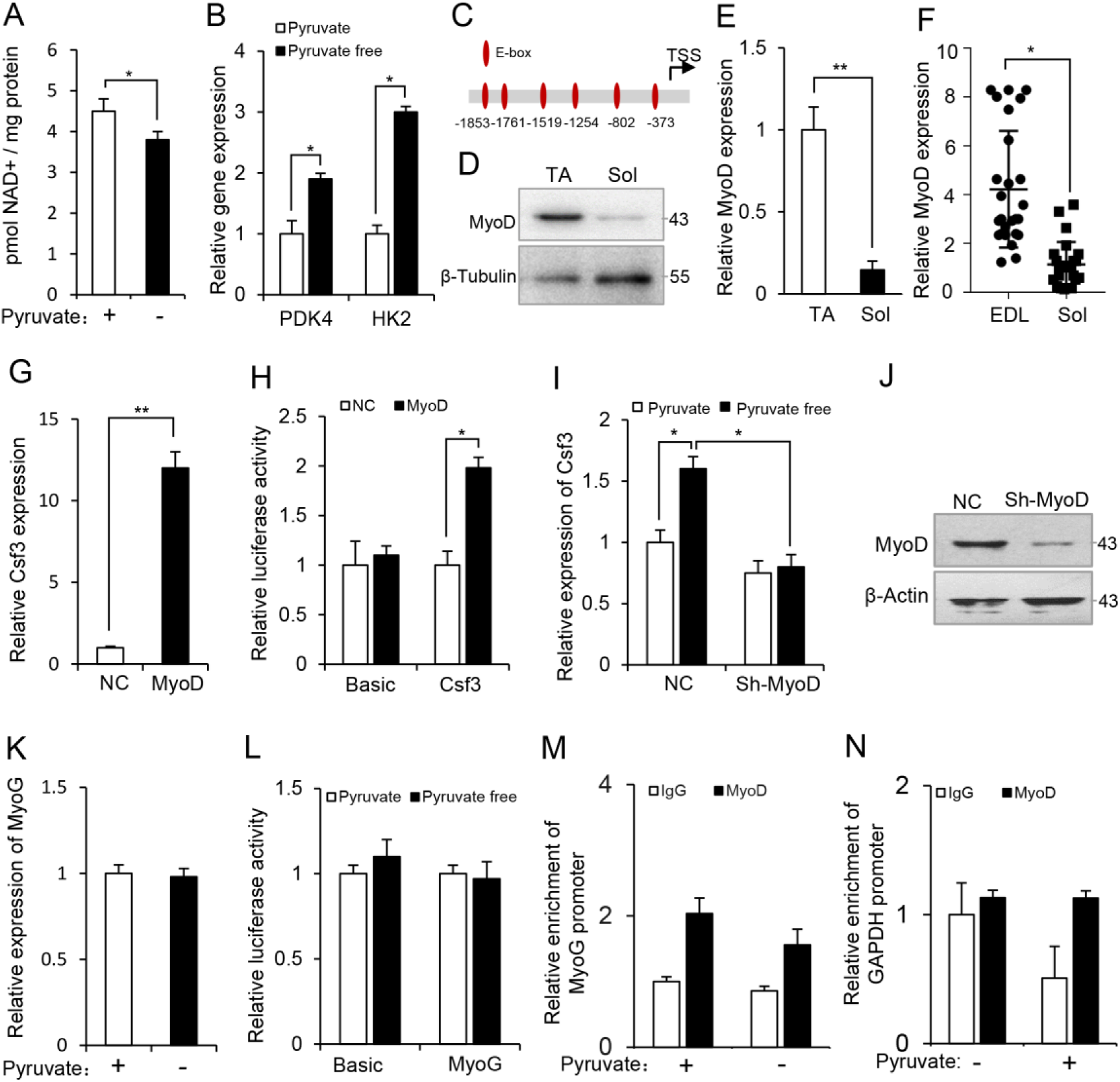
MyoD metabolically regulates *Csf3* gene expression in skeletal muscle cells. A NAD+ levels in C2C12 myotubes cultured in medium with or without pyruvate for 24 hr. Data were obtained from 3 independent experiments. *p< 0.05. Unpaired two-sided t-test. B Relative expression of *PDK4* and *HK2* genes in the C2C12 myotubes cultured in medium with or without pyruvate for 24 hr. Data were obtained from 3 independent experiments. *p< 0.05. Unpaired two-sided t-test. C E-boxes (CANNTG) within the 2-kb sequence upstream of the mouse *Csf3* gene. D Protein levels of MyoD in TA and Sol muscles, as determined by Western blotting. β-Tubulin served as a loading control. E mRNA levels of *MyoD* in TA and Sol muscles, as determined by qRT-PCR. *β-Actin* served as an internal control. n=4. *p< 0.05. Unpaired two-sided t-test. F Relative expression of *MyoD* gene in isolated single fiber from Sol and EDL muscles was determined by qRT-PCR. Each dot represented one single myofiber. n=25 in Sol group. n=24 in EDL group. *p< 0.05. Unpaired two-sided t-test. G Relative expression of *Csf3* in C2C12 myotubes overexpressing MyoD. Empty vector as negative control (NC). Data were obtained from 3 independent experiments. *p< 0.05. Unpaired two-sided t-test. H Relative promoter activities of the 2-kb sequence upstream of *Csf3* in fibroblasts with transient ectopic expression of *MyoD*. Transfection with empty vector served as a negative control (NC). Data were obtained from 3 independent experiments. *p< 0.05. Unpaired two-sided t-test. I Relative expression of *Csf3* in MyoD-knockdown C2C12 myotubes cultured in medium with or without pyruvate. Data were obtained from 3 independent experiments. *p< 0.05. 2-way ANOVA. J MyoD protein levels in the MyoD-knockdown C2C12 myotubes. β-Actin served as a loading control. K Relative expressions of *MyoG* in C2C12 myotubes cultured in medium with or without pyruvate for 24 hr. L Relative promoter activities of *MyoG* in C2C12 myotubes cultured in basal or pyruvate-free medium for 24 hr. M,N ChIP-qPCR were performed using chromatin from myotube cultured in presence or absence of pyruvate. Chromatin was immunoprecipitated using antibodies against MyoD. The immunoprecipitated DNA was amplified using primers for *MyoG* (M) and *GAPDH* (N) genes promoter.

**Fig EV6.**
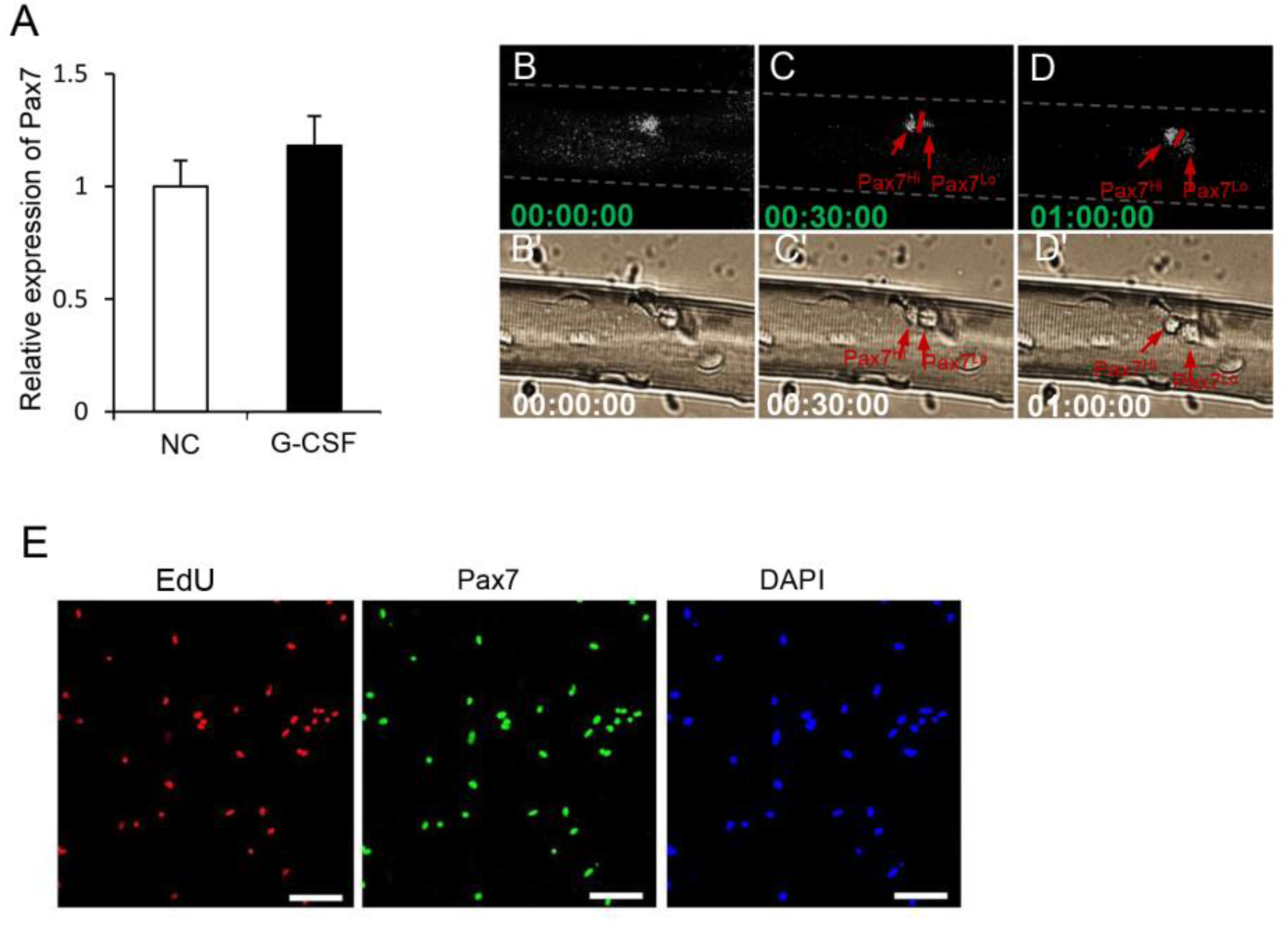
G-CSF promotes asymmetric division of Pax7 SCs. A Relative expression of *Pax7* in FACS-resolved Pax7 SCs cultured in GM containing G-CSF for 24 hr, as determined by qRT-PCR. Data were obtained from 3 independent experiments. B-D Time-lapse images were used to trace the division of Pax7 SCs on single fibers isolated from *Pax7-nGFP* mice, n=4. E Representative EdU staining in FACS-resolved Pax7 SCs from *Pax7-nGFP* mice at T1. Scale bar represents 40 µm.

**Fig EV7.**
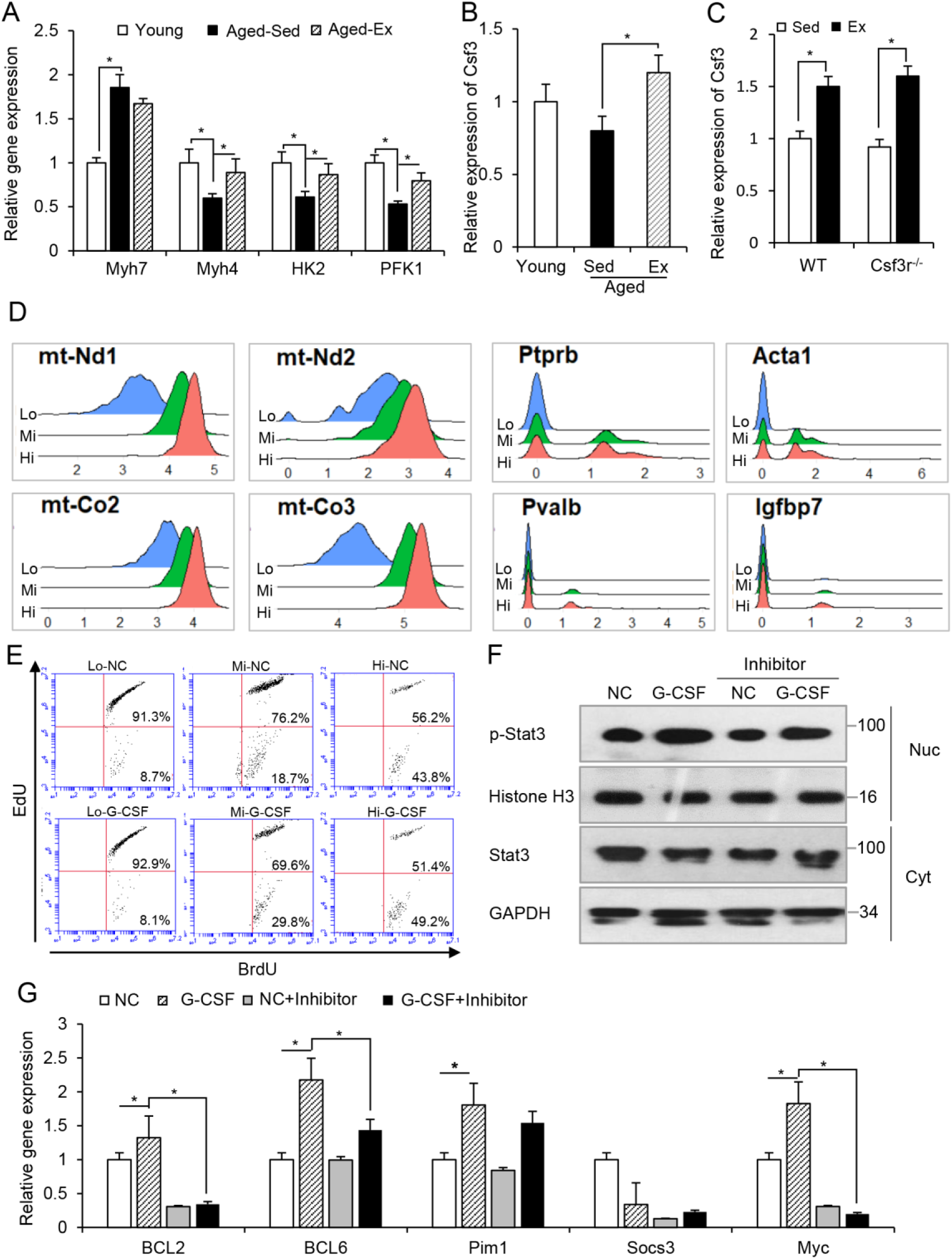
G-CSF replenished Pax7^Hi^ cells by promoting asymmetric division of Pax7^Mi^ cells. A Relative expression of *Myh7*, *Myh4*, *HK2,* and *PFK1* in TA muscles from young (Young), aged exercised (Aged-Ex), and aged sedentary (Aged-Sed) Pax7-nGFP mice. *n* = 5. *p< 0.05. 1-way ANOVA. B Relative expression of *Csf3* in TA muscles from young (Young), aged exercised (Aged-Ex), and aged sedentary (Aged-Sed) Pax7-nGFP mice. *n* = 5. *p< 0.05. 1-way ANOVA. C Relative expression of *Csf3* gene in the TA muscles of *Pax7-nGFP;Csf3r^-/-^* or *Pax7-nGFP* WT mice subjected to exercise (Ex), as determined by qRT-PCR. *n* = 4. *p< 0.05. Unpaired two-sided t-test. D Representative genes in Pax7^Hi^, Pax7^Mi^ and Pax7^Lo^ cells were visualized by Ridge plots. E Flow cytometric analysis of the three subpopulations of Pax7 SCs treated with G-CSF for 24 hr and subjected to EdU and BrdU immunostaining. Numbers in corners represented percentages (%) of the cells. The three subpopulations of Pax7 SCs were sorted from Pax7-nGFP mice and pulse-labeled with EdU and BrdU. F Protein levels of Stat3 and p-Stat3 in Pax7 SCs treated with G-CSF in the presence or absence of the Stat3 inhibitor, 5,15 DPP, as determined by Western blotting. GAPDH and histone H3 served as loading controls for the cytoplasmic and nuclear fractions, respectively. G Relative expression of *BCL2*, *BCL6*, *Pim1*, *Socs3*, and *Myc* in Pax7 SCs treated with G-CSF in the presence or absence of 5,15 DPP, as determined by qRT-PCR. Values were presented as the mean ± SEM. Data from three independent experiments. *p< 0.05. 2-way ANOVA.

